# Dynamic modelling of human neural crest development using a bioengineered stem cell organoid system

**DOI:** 10.64898/2026.05.04.721958

**Authors:** Carmen Moreno-Gonzalez, Dylan Cameron, Marcela Marques Moreno, Jade Desjardins, Taylor F. Minckley, Matthew C.D. Bailey, Cathleen Hagemann, Shail U. Bhatt, Anestis Tsakiridis, Andrea Serio, Karen J. Liu

**Author notes:** Equal co-corresponding authors:.

## Abstract

The neural crest (NC) is a transient stem cell population which migrates throughout the developing embryo to contribute to diverse tissues dependent on axial origin. For example, cranial NC can give rise to bone and cartilage, while more posterior NC populations give rise to peripheral nervous system and neuroendocrine tissues. Perturbations in neural crest development can lead to severe congenital anomalies and cancers, with over 700 neurocristopathies reported. In humans, early NC development remains poorly understood due to the inaccessibility of tissue samples, thus necessitating the development of *in vitro* models. Currently, a limited number of NC organoid protocols are available, but these mainly focus on cranial NC and lack relevant tissue architecture. Here, we describe a novel bioengineered pipeline to derive human pluripotent stem cell (hPSC)-derived neuroepithelial organoids, “neurocrestoids” featuring physiologically-relevant tissue architecture. We show that neurocrestoids recapitulate the dynamics of induction, delamination, and migration of human neural crest cells (NCCs), and can be directly compared to murine NC explants for cross-species validation. Organoids express an array of *HOX* genes indicating the successful generation of cranial, vagal and trunk NCCs. Moreover, we have integrated our neurocrestoids with a customised micropatterned substrate suitable for live visualisation and guided separation of SOX10-positive migratory human NCCs. Our “NCC migration on-chip” are reproducible across multiple hPSC lines and should be scalable for future diagnostic and therapeutic applications, significantly improving our ability to study human NC pathologies.

## Introduction

Neural crest cells (NCCs) are migratory embryonic stem cells contributing to most organs. They are uniquely plastic, capable of generating over 30 cell types dependent on their axial identity. NCCs are specified at the neural plate border (NPB) and undergo epithelial-to-mesenchymal transition (EMT) to delaminate from the dorsal neural tube and migrate long distances toward target tissues^1,2^. NCCs arise along the entire anteroposterior body axis, including cranial, vagal, trunk and sacral lineages, each having a precise developmental tempo, specific regulatory elements, cell behaviours and differentiation potential^3,4^.

Previous *in vivo* experiments in model systems demonstrate that NCCs arise from the NPB during gastrulation, requiring a complex interplay of Wnt, FGF, BMP and Notch signals from surrounding tissues. These studies also reveal fundamental species-specific differences in gene expression and developmental timing during NC formation^4–6^. However, the degree to which human NC behaviours align with established animal models remains poorly understood due to limited access to human embryonic tissue samples and ethical constraints on performing live and interventional studies. This is particularly challenging in the context of NC development, where their developmental timing coincides with an early phase of human embryogenesis, often referred to as the “black box”, a period between the current legal limit for *in vitro* embryo culture^7^ and the availability of the earliest embryonic samples.

Recently, transcriptomic profiling of human embryos revealed that the first anterior NCC populations are induced as early as Carnegie stage (CS) 9 (week 3 post-conception), coinciding with the conclusion of gastrulation^8,9^. From CS10, cranial NCCs become evident and delaminate before neural tube closure, and, by CS12, cardiac NCCs arise and begin migration^10^. To our knowledge, only a few studies have investigated the relationship between the spatial distribution and the expression of NC-specific genes during human development, demonstrating the presence of *bona fide* markers such as Zic1, SOX9, SOX10, p75NTR, and AP2α^11,12^. Nonetheless, human NC development remains poorly understood. Notably, NC defects result in a broad spectrum of more than 700 disorders named neurocristopathies, which affect a high proportion of live births^13^, creating a pressing need for better human *in vitro* models.

In recent years, human pluripotent stem cell (hPSC)-derived three-dimensional culture systems have emerged as attractive approaches to studying human embryogenesis, with a long-term goal of linking patient genotypes to mechanisms of disease. By integrating multiple cell types within physiologically relevant tissue architecture, these enable the investigation of genetic and environmental influences (e.g.^14-16^). Reflecting this broader trend, there has been a growing emphasis on developing organoid models in the NC field. Several studies described the generation of human embryonic stem cell (hESC)-derived neural spheres capable of producing migrating NCCs upon plating^17–19^. Others have shown the generation of NCCs in 3D using cranial NC organoids, neural tube models or combined neuroectodermal and mesodermal assembloids^20–22^. While these 3D-cultures can mimic certain *in vivo* events, they rarely represent the full range of NC development or tissue complexity, and are mostly restricted to cranial NC.

To overcome these limitations, our goal was to develop a robust human-specific *in vitro* model system capable of generating the full range of NCCs, recapitulating the 3D architecture and spatiotemporal dynamics of neural crest induction, EMT and delamination, followed by isolation of migratory NCCs. Moreover, we aimed to obtain minimal variability across multiple batches and lines. Learning from previous organoid-based protocols^23–25^, we decided to address reproducibility and scalability inherent to 3D self-organising cultures (e.g. 3D aggregate size and diameter distribution), using microfabrication techniques to generate customised culture devices. Specifically, biocompatible microwells have previously been employed to control and standardise the dimensions and shape of starting spheroid cultures^26,27^.

In this study, we describe the generation of a bioengineered neuromesodermal organoid platform capable of robustly producing an array of NC subtypes within a physiologically relevant 3D tissue architecture. This approach can then be combined with micropatterned polydimethylsiloxane (PDMS) substrates to selectively guide and visualise NCC migration. The starting material for our protocol is the generation of large-scale batches of homogeneously generated EBs using custom-made PDMS microwells. To differentiate organoids, we adapted established principles from previously published 2D protocols, which produce NCCs via neuromesodermal progenitors (NMPs)^28,29^. Combining these, we obtained organoids with distinctive epithelial structures containing major embryonic cell types, producing an array of NC subtypes including cranial, vagal and trunk NCCs. These organoids faithfully recapitulate the temporal progression of key milestones in NC development. Importantly, neurocrestoids reveal an organoid architecture resembling embryonic patterns of key transcription factors, with SOX9 localised to pre-migratory NCCs in epithelial regions, and SOX10 marking migratory NCCs, which we compare to *in vivo* distribution using mouse embryos.

Next, we demonstrated that neurocrestoids generate NCCs that migrate when plated, which we directly compared to primary mouse NCCs for cross-species validation. We demonstrated that combining our neurocrestoids with customised microgroove devices allows us to guide, segregate, and directly visualise human NCCs, thus, generating a novel, human-specific “NCC migration-on-chip” platform. Using this model, we uncovered a strong preference of NCCs towards laterally confined microchannels, allowing us to isolate NCCs from other confounding populations.

In conclusion, we have created a robust human stem-cell-derived organoid system capable of recapitulating NC development along the anteroposterior axis and coupled it with micropatterned PDMS substrates to guide NC migration. We anticipate that this platform will advance our understanding of human NC development, and, given its robustness and scalability, it will provide an unprecedented tool for disease modelling and drug discovery.

## Results

### Human stem cell-based pipeline using custom-engineered microwells yields neuroepithelial organoids capable of generating NCCs in 3D

We first generated hPSC-derived embryoid bodies using custom-made PDMS microwells. Previous methods for generating neuroectodermal organoids relied on spontaneous aggregation^17,18^, which yield irregular aggregate sizes that can impact downstream differentiation outcomes, or on commercially available systems, which generally lack the possibility of customisation^21,22^. Using a pipeline we recently developed^26^, we were able to customise PDMS microwell devices (**Fig. S1)** and select an optimal size (400x400x250µm) for our experiments, designed to support reproducible and scalable aggregation of EBs from an hPSC suspension. To optimise our PDMS microwells, we started from previously tested designs and increased the well-depth to avoid contiguous EB fusion, while maintaining a depth that minimised air bubble formation (**Fig. 1A-C, S1A-C**). EBs were then transferred to a rotatory culture until day 8, when we collected samples or plated the organoids for downstream applications (**Fig. 1A**). Our system reproducibly generates consistently-sized EBs (median diameter 243µm) in an effective and high-throughput manner (**Fig. 1D**).

**Fig. 1.**
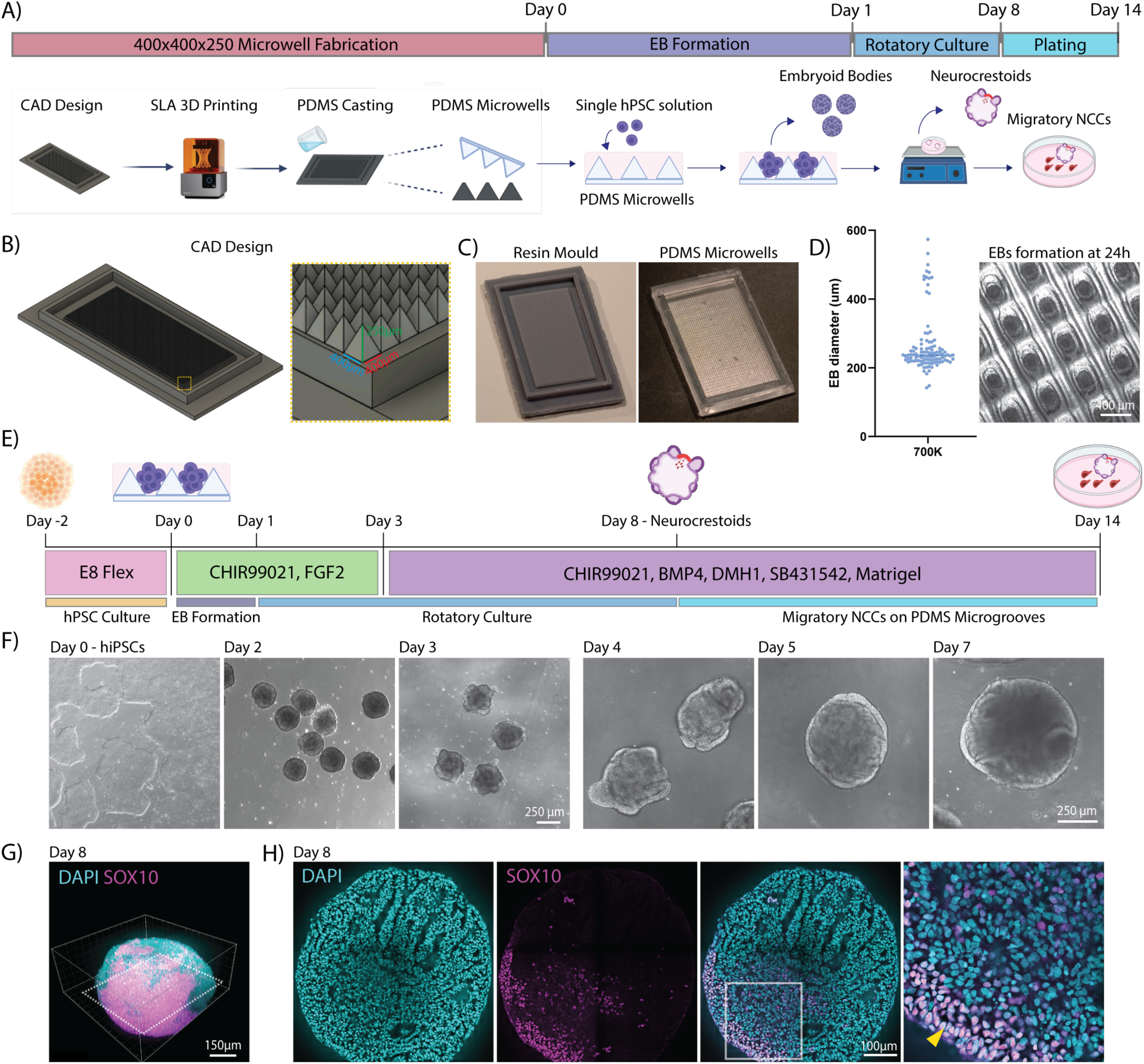
– Development of a bioengineered pipeline for the generation of neuroepithelial organoids capable of producing NCCs in 3D and upon plating using custom-made bioengineered microwells. A) Schematic overview of the pipeline for generating neurocrestoids using custom-fabricated PDMS microwells. Prior to the start of the differentiation protocol, PDMS microwells are fabricated from custom-made resin moulds to aggregate hPSCs into spheroids as previously described^26^. B) Overview of the CAD design for 400x400x250µm inverted pyramidal-shaped PDMS microwells. C) Images of the custom-made resin mould and resulting PDMS microwells. D) Graph indicating the sphere diameter (median 243µm), accompanied by an image of spheroids in PDMS microwells at 24 hours after seeding. Points represent single organoids (n=105). E) Schematic depicting the protocol to generate hPSC-derived neuroepithelial organoids, termed neurocrestoids, capable of producing NCCs in 3D. F) Representative brightfield images during the first 7 days of the differentiation process. G) 3D rendering from a whole-mount immunostained suspension organoid presenting SOX10-positive regions, demonstrating its ability to generate NCCs in 3D. H) A representative optical slice from the organoid in F) showing internal epithelial structures from which SOX10-positive cells emanate (yellow arrowhead). Scale bar = 100µm.

To differentiate EBs toward posterior NC, we adapted a previously described 2D protocol for the derivation of trunk NCCs from hPSCs, which is achieved via a *SOX2/TBXT-*positive NMP-like state (**Fig. 1E**)^29,30^. We hypothesised that an early pulse of FGF2 and CHIR99021 (CHIR), which activates canonical Wnt signalling, would bias the EBs towards an NMP population capable of differentiating into NCCs, as previously shown in 2D^29,31^. After day 3, organoids were treated with BMP4 and DMHI, a BMP inhibitor, as part of a “top-down” approach to finely tune the activation of BMP signalling^32^. This medium was complemented with a TGFβ inhibitor and sustained Wnt activation to allow for NC specification. In addition, we introduced Matrigel to support the formation of complex architectures such as epithelial structures, which were visible from day 3 to 4 (**Fig. 1F**).

Whole-mount immunofluorescence staining of neurocrestoids at day 8 shows confined SOX10-positive regions, indicating the production of NCCs (**Fig. 1G**). Optical slices of these organoids reveal the presence of epithelium-like structures with adjacent SOX10-positive cells reminiscent of delaminating NC (**Fig. 1H**). Similar results were observed with three additional hPSC lines across two independent laboratories (**Fig. S1D**).

Early Wnt activation has been extensively reported to play pivotal roles in NC development and determination of axial identity, both *in vivo* and *in vitro*^29,33–36^. In line with this, modulating the levels of canonical Wnt activation during the first 3 days via CHIR addition was a crucial determinant of tissue composition, architecture and NC generation. Lower CHIR dosages resulted in a higher proportion of neuroepithelia, upregulation of NMP and NC markers and 3D-NCC generation at day 8. Neurocrestoids progressively lose their NMP and NC signature as we increased CHIR concentrations, as shown by the downregulation of *SOX2, CDX2, TBXT* and *SOX10* markers (**Fig. S1E and S1H**).

### Neurocrestoids are recreate the dynamics of NC development

Having established a protocol to generate organoids capable of producing NCCs, we tested their ability to mimic the developmental events in a temporally coherent manner as well as their capacity to generate axial diversity in the NCCs. We isolated neurocrestoids at sequential stages: day 0, 3, 6, and 8, to capture key developmental timepoints: the emergence of axial progenitors (day 3), an intermediate differentiation stage (day 6) and the generation of NCCs (day 8), as well as isolating NCCs obtained by dissociation and sorting day 8 organoids using p75NTR-magnetic beads (N=3) (**Fig. 2A**).

**Fig. 2.**
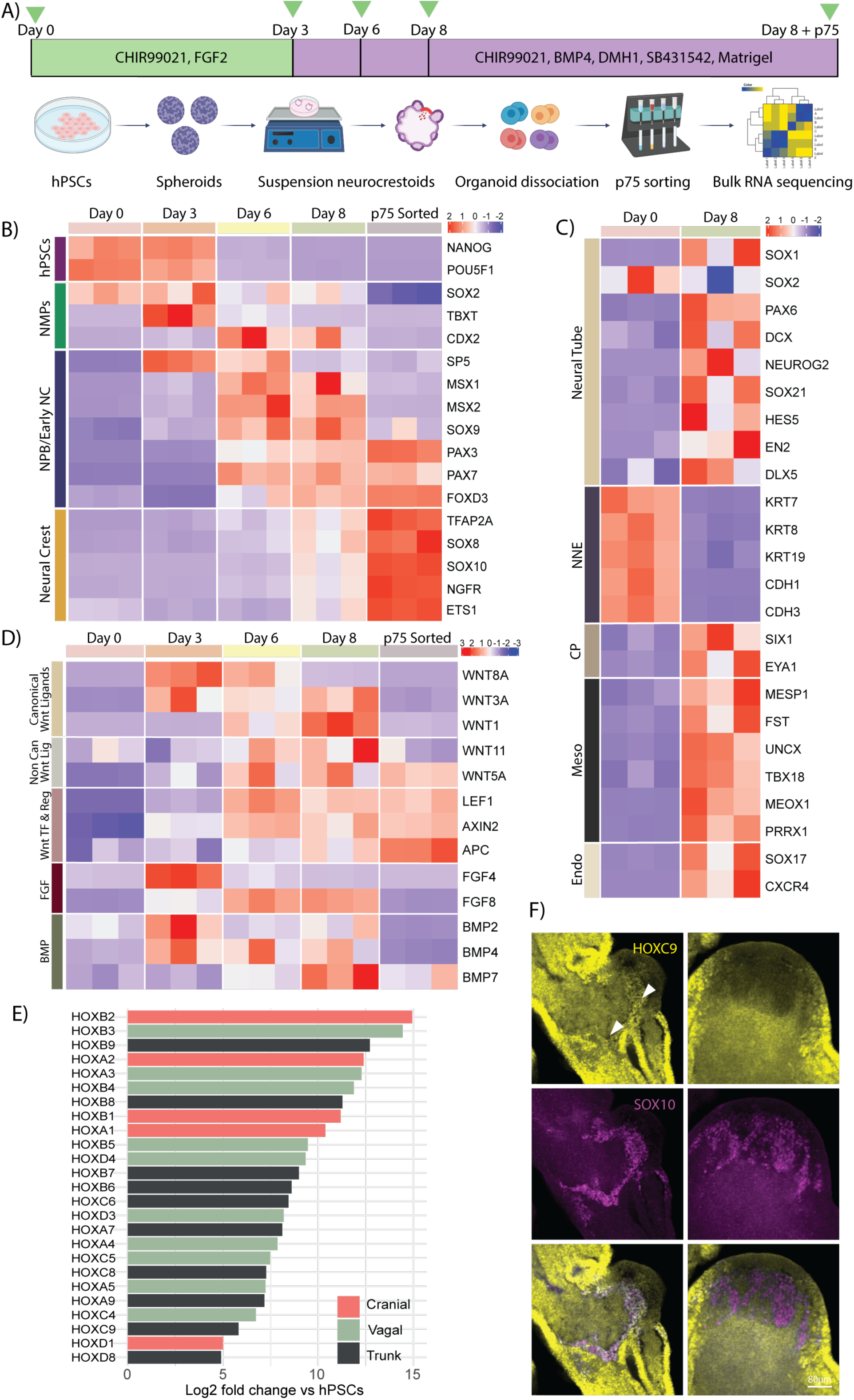
- Neurocrestoids are neuromesodermal organoids which recapitulate transcriptional signatures hallmark of NC development to generate NCCs from various axial identities. A) Schematic depicting the pipeline of the RNA sequencing experiment. Green arrows represent timepoints when organoids were sampled. Note that at day 8, neurocrestoids are dissociated into single cells, and NCCs are selected using p75NTR magnetic beads and added to the sample pool for transcriptomic analysis. B) Heatmap of differentially expressed genes (padj<0.05; |log2FC|>1; D0 vs p75) relating to pluripotent, NMP, NPB/Early NC, and NC populations over time. C) Heatmap of differentially expressed genes (padj<0.05; |log2FC|>1; D0 vs D8) relating to NT, NNE, CP, mesoderm, endoderm and NC populations in day 8 neurocrestoids. NNE: Non-neural ectoderm, CP: Cranial Placodes, Meso: Mesoderm, Endo: Endoderm, NC: Neural Crest. D) Heatmap of differentially expressed genes relating to Wnt canonical pathway ligands, Wnt non-canonical pathway ligands, Wnt canonical pathway effectors and regulators, FGF and BMP ligands over time. E) Bar chart depicting the change in expression (log2 fold change) in the NCC condition (p75 sorted) for all statistically significantly expressed Hox genes versus day 0. Different shades of green mark Hox genes relating to cranial, vagal and trunk axial positions. Hox genes are in descending order by log2 fold change in expression versus day 0. F) Example optical slices of 2 different organoids showcasing the ability of neurocrestoids in generating different subpopulations of NCCs at day 8. White arrowheads indicate SOX10-positive areas.

Principal component analysis (PCA) showed clustering of samples according to differentiation stage, with minimal variation between technical replicates (**Fig. S2A**). To prove the successful isolation of NCCs, we investigated the normalised counts of p75 transcripts and compared them to SOX10 and PAX6 counts. We detected a clear enrichment of SOX10 in the p75-sorted sample, as opposed to PAX6, a CNS marker, corroborating the suitability of p75 beads to isolate NCCs from dissociated neurocrestoids (**Fig. S2B**).

Differential expression analysis of our RNA-seq dataset confirmed progressive onset of developmental milestones. First, upregulation of NMP markers (*SOX2/TBXT)* was seen at earlier timepoints (days 3 to 6), followed by the expression of NPB and early NC markers such as *SP5, MSX1/2, PAX3, and PAX7* (day 6), before the emergence of *bona fide* NC genes (*FOXD3, SOX9, TFAP2A, SOX10, P75NTR, ETS1*) at day 8. Moreover, the transcriptomic profile of p75-sorted condition demonstrates the ability of neurocrestoids to generate NCCs that express relevant pre- and pro-migratory NCC markers such as (*PAX3, PAX7, FOXD3, TFAP2A, SOX10, P75NTR, ETS1*) (**Fig. 2B**).

Supporting the heterogeneity seen *in situ* (**Fig. 1G-H**) and reminiscent of *in vivo* conditions, day 8 neurocrestoids expressed transcriptional signatures from all embryonic lineages, including neural ectoderm, mesoderm and cranial placode markers (**Fig. 2C**). Neurocrestoids show a total of 4058 and 4013 genes up- and downregulated versus day 0, respectively (padj<0.05; |log2FC|>1; **Fig. S2C** and **S3**). Gene ontology (GO) enrichment analysis reveals a preference for genes involved in organ development and anteroposterior patterning, highlighting the ability of neurocrestoids to mimic the complex embryonic environment. Notably, GO analysis for the p75-sorted condition, which enriches for NCC, confirms acquisition of anteroposterior patterning genes (**Fig. S3A-B**). Interestingly, when compared to day 8, the GO analysis shows an enrichment of terms related to neuronal development and myelination in the P75-sorted condition (**Fig. S3C**). Concurrently, we observed an upregulation of genes related to sympathetic neurons (*ELAVL3/4, DLL4, TH, PHOX2B*), Schwann cell precursors (*DLX1/2, MPZ, PLP1, S100B*), melanocytes (*FOXD3, MLANA, SOX10*) and sensory neurons (*ONECUT2, NEUROG1, ISL1, NTRK1*), but not of smooth muscle, chondrocyte, chromaffin cell or osteoblast markers at day 8 neurocrestoids (**Fig. S2D**).

Differential expression analysis also demonstrates the presence in a time-coherent manner of specific factors of the Wnt, Fgf and BMP signalling pathways, crucial for NPB specification and NC induction *in vivo*^37–40^. From day 3, we note an upregulation of ligands and effectors of the Wnt canonical pathway: the mesoderm-produced *WNT8A*, and *WNT3A* and *WNT1*, which are typically expressed by the NPB and NC. From day 6 onwards, we also detect the presence of non-canonical Wnt ligands *WNT5A* and *WNT11*, which are key for NC induction and migration in animal models. Additionally, essential ligands of the FGF and BMP pathways are also upregulated in a temporally coherent manner (*FGF4*, *FGF8*, *BMP2, BMP4)*. In p75-positive cells, we observe expression of *LEF1* and the negative feedback regulators *AXIN2* and *APC*, indicating initial activation of the canonical Wnt pathway and subsequent inhibition of Wnts, which is known to be required as NC begin to migrate (**Fig. 2D**). Taken together, these results demonstrate the ability of neurocrestoids to recapitulate NC transcriptional signatures and reflect the complex signalling microenvironment from which NCCs arise, highlighting the integration of multiple cell types within our system.

### Neurocrestoids generate an array of NCCs along the anteroposterior axis, including cranial, vagal and trunk NC

In the embryo, the NC can be subdivided into four main functional domains depending on their position along the anteroposterior axis: cranial, vagal, trunk and sacral NC. Currently available NC organoid protocols have not yet demonstrated the generation of NCCs in 3D beyond the cranial compartment, leaving a gap for novel models that can recapitulate posterior NC development^17,18,21,22^. These sub-lineages have distinct developmental timing, regulatory elements, migratory behaviours and final derivatives^41^, with their axial identity indicated by the progressive expression of *HOX* genes. When we examined the p75-sorted condition, we found an array of Hox genes ranging from *HOXA1 to HOXC9/HOXD8*, suggesting the emergence of posterior cranial, vagal and trunk NCCs in our organoids (**Fig. 2E**). To validate the presence of the more posterior NC subtypes, we performed immunocytochemistry assays of whole-mount organoids at day 8, labelling for HOXC9 and SOX10 (**Fig. 2F**). Day 8 neurocrestoids include both HOXC9-positive and negative NCC populations, corroborating the generation both anterior and posterior NC, including thoracic trunk NCCs in 3D, independently of microfluidic gradient control (**Fig. 2F)**. Taken together, these results demonstrate that our protocol produces multiple axial subtypes in 3D, including cranial, vagal and trunk NC.

### Neurocrestoids recapitulate NC delamination and EMT in a physiologically relevant three-dimensional environment

*In vivo*, NCCs acquire mesenchymal characteristics that allow delamination, indicated by a defined switch in expression of cadherin isoforms and onset of EMT markers^42,43^. Differential expression analysis of our RNA-seq dataset reveals a switch from *CDH1* (*E-cadherin*) expression to *CDH2* (*N-cadherin*) between days 3 and 6, reflecting the dynamics of NC and mesoderm development. Type II cadherins were also expressed similarly, with *CDH7* evident by day 3, and *CDH6* and *CDH11* from day 6, which are typically expressed in migratory NCCs. Other EMT proteins were also seen in a temporally coherent manner (*TWIST1, SNAI1/2, ZEB1/2*) (**Fig. 3A**). Interestingly, we consistently observed SOX9-positive cells within the epithelium and adjacent SOX10-positive regions (N=3), which expanded towards the centre of the organoids. At the interface between these two domains, we could detect a layer of double-positive SOX9/SOX10 NCCs, suggesting a transition between the two transcription factors (**Fig. 3B**).

**Fig. 3.**
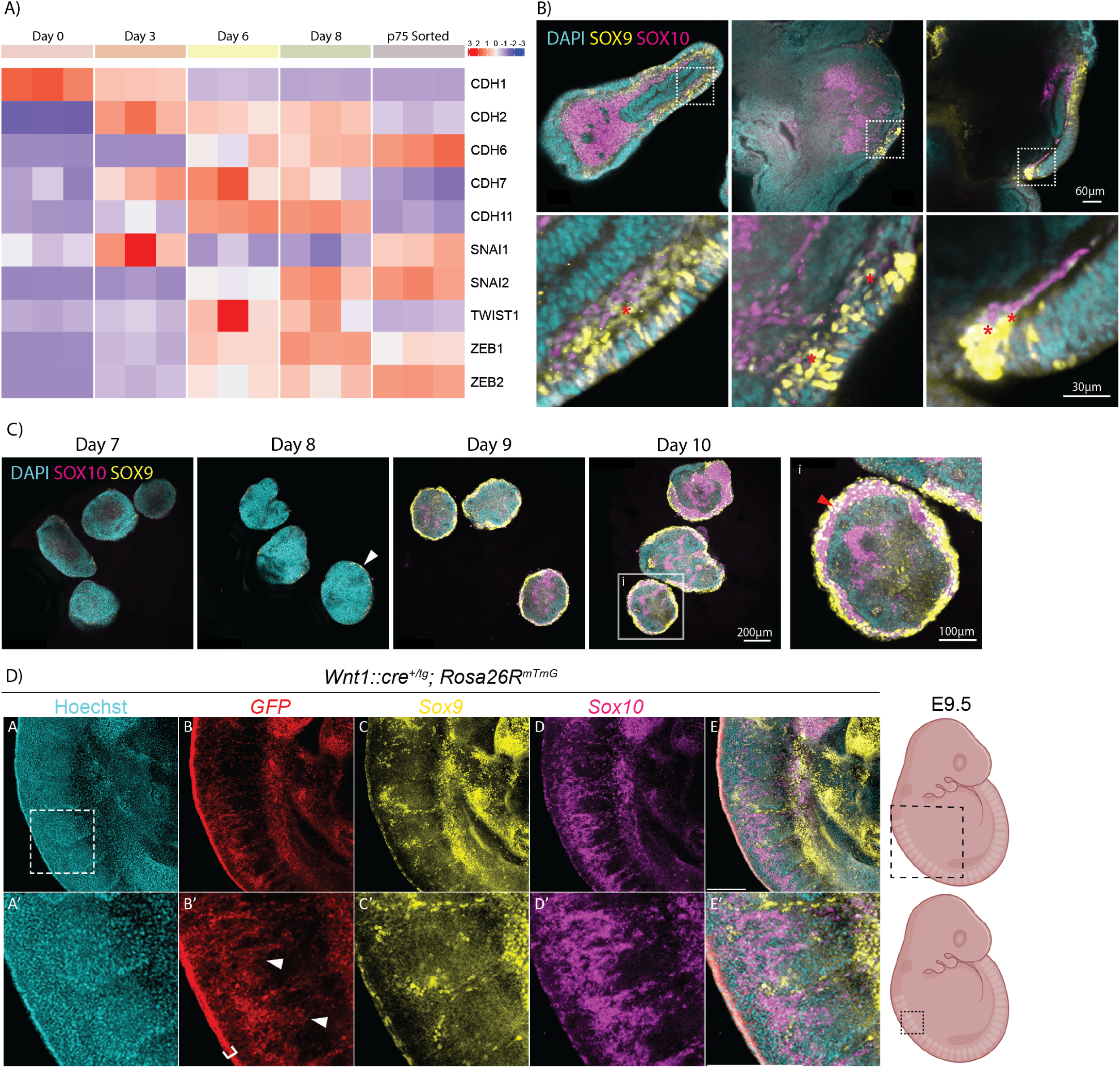
– Neurocrestoids recapitulate NC delamination in a physiologically relevant tissue architecture and showcase a SOX9/SOX10 transition reminiscent of in vivo conditions. A) Heatmap of differentially expressed EMT genes (D8 vs D0) through the neurocrestoid differentiation process, including Cadherin isoforms. B) Representative Z-stacks of day 8 neurocrestoids from 3 independent neurocrestoid experimental replicates, along with zoomed-in images of the areas indicated by the white dashed squares. Note the presence of SOX9-positive cells in epithelial structures immediately adjacent to SOX10 populations and separated by a double-positive cell layer (red asterisks). C) Representative images of neurocrestoids through days 7 to 10 of the differentiation protocol, immunolabelled for SOX9, SOX10 and nuclei (DAPI). SOX9 populations appear at day 8 (white arrowhead). Accompanying is a zoomed-in image of a neurocrestoid at day 10 (i), which presents SOX9-positive cells in the edges, a double-positive layer of cells (red arrowhead) and SOX10 populations towards the organoid core. D) Representative maximum intensity projection of an HCR on an E9.5 Wnt1::cre^+/tg^; Rosa26R^mTmG^ mouse embryo labelling GFP (NCCs), Sox9, Sox10 and nuclei (Hoechst) in the trunk regions (somites 1 to 10).

Previous *in vivo* studies have shown the ability of *Sox9* to act as an upstream regulator of *Sox10* expression in NCCs^44,45^. To investigate the order and developmental timing of the two transcription factors, we sought to investigate the appearance of SOX9/SOX10-positive structures within our organoids over time. Whole-mount immunocytochemistry images of day 7 to 10 organoids show that SOX9 becomes apparent within epithelia at the organoid edges at day 8, at a time when the SOX10 signal is yet undetected. By day 9, the presence of SOX9, SOX10 and double-positive regions can be seen, revealing that this transition happens within 24 hours (**Fig. 3C**). These data suggest that SOX9 is established in epithelial NCCs prior to SOX10 expression in mesenchymal NC in human NCCs.

To relate our previous findings to *in vivo* conditions, we compared the spatial arrangement of *Sox9* and *Sox10* in NCCs from mouse embryos via *in situ* hybridisation chain reaction (HCR), using transgenic mice carrying an NC-specific reporter (*Wnt1::cre^+/tg^; Rosa26R^mTmG^*) (N=4). In these animals, GFP-positive migratory NCCs are visualised at embryonic day 9.5 (E9.5) in the trunk region. As anticipated, *Sox9* mRNA was observed predominantly in premigratory NCCs of the dorsal neural tube, while the majority of delaminating and migrating GFP-positive NCCs expressed *Sox10,* extending ventrally towards the spinal nerves (**Fig. 3D**). Overall, these data demonstrate the similarities between the spatial and functional organisation of two hallmark TFs in NC development, SOX9 and SOX10, in mouse and human organoids, highlighting the power of neurocrestoids to recapitulate key events of NC development in association with physiologically relevant architecture, providing an unprecedented opportunity to study human NC delamination.

### Plated neurocrestoids generate SOX10-positive migratory NCCs with unique and distinct cellular and nuclear characteristics

Previous studies have shown the ability of neural spheroids to generate migratory cranial NC populations when plated in cell culture dishes^17,18^, which represents an attractive approach to studying NC migration. Similarly, we plated day 8 neurocrestoids on Matrigel-coated plates, which resulted in migratory cell populations that extended radially over the area of the well (**Fig. 4A** **and S4A**). We cultured plated neurocrestoids for an additional 6 days and performed immunofluorescence assays at day 14 to identify SOX10-positive populations amongst the migratory cell pool (mean 62% SOX10-positive out of the total migrated area across experiments and hPSC lines, N=3) (**Fig. 4B-C** **and S4A**).

**Fig. 4.**
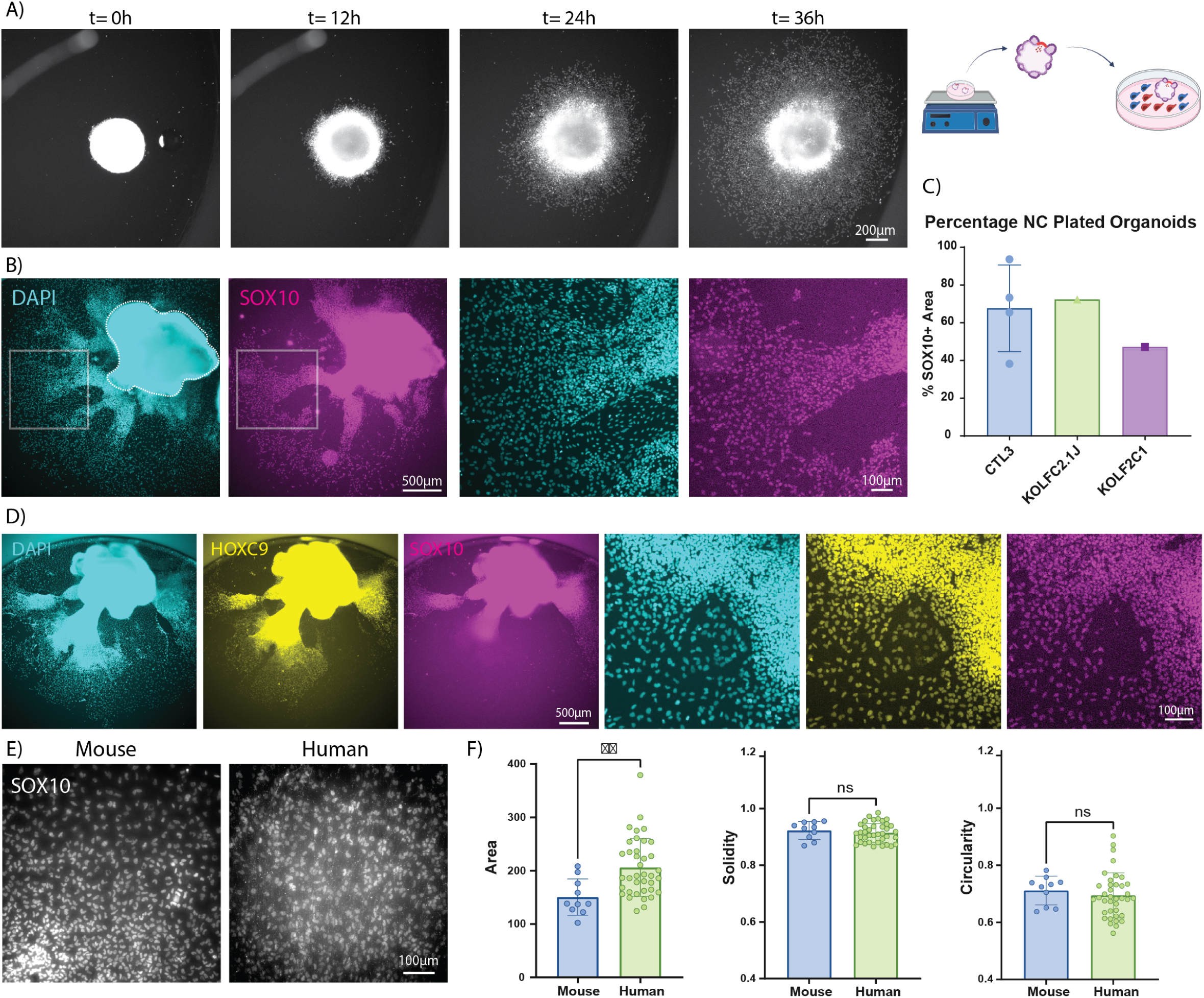
– Plated neurocrestoids generate SOX10-positive migratory NCCs and non-SOX10-positive cell populations with distinct cellular characteristics, which restrict NCC migration. A) Time-lapse images of a live plated neurocrestoid with nuclei stained with SPY650-DNA. B) Representative fluorescence images at day 14 of plated neurocrestoids immunostained for SOX10 and nuclei (DAPI). C) Bar chart depicting percentage of SOX10 positive cells amongst the total nuclei (DAPI) in plated neurocrestoids at day 14 across experiments from 3 independent hPSC lines (mean ± S.D). Each point represents the average across n=5-18 organoids per individual experiment. F) Neurocrestoids plated at day 14 and stained against HOXC9, SOX10 and nuclei (DAPI). E) Representative immunofluorescence images of mouse trunk explant and human neurocrestoid-derived NCCs stained for SOX10. F) Bar charts indicating the area, circularity and solidity values for neurocrestoid-derived human and mouse trunk NC nuclei (mean ± S.D). Points represent the average value per field of view. For the human datasets, a total of 38 datapoints are plotted representing the average value per field of view, 5 images per sample, 3-5 organoids per cell line from 3 hiPSC lines (Control 3, KOLFC21 and KOLF2.1J) in 3 experimental blocks. The mouse dataset consists of the average of 10 cells per image, 10 images per explant in 3 mouse trunk NC explants cultured for 48 to 72 hours in 2 experimental blocks. Non-parametric Mann-Whitney test to test for significance between conditions, **P<0.01, ns: non-significant.

As seen with day 8 suspension organoids, early Wnt activation was crucial for the amount of NCCs emigrating from plated neurocrestoids at day 14. By measuring the area covered by SOX10-positive cells, we observed that CHIR dosage was positively correlated with a reduction in migrating NCCs. Interestingly, intermediate CHIR dosages also produced migratory NCCs in plated neurocrestoids, which allowed us to define 3 distinct groups based on organoid architecture and their ability to generate NCCs: “low”, “medium” and “high” CHIR conditions, with only the first consistently yielding NCCs in suspension and upon plating (**Fig. S1F, S1H and data not shown**).

Concurrent with RNA-seq results, immunocytochemistry labelling of HOXC9 and SOX10 confirmed the generation of migratory trunk NCCs from plated neurocrestoids (N=3) (**Fig. 4D**). Higher magnification immunofluorescence images also revealed distinct actin localisation in SOX10-positive versus SOX10-negative cell types, with NCCs present in high cell density areas, and showing disorganised and superimposing actin filaments, with bean-shaped nuclei (**Fig. S4D**). During cell migration, the nucleus undergoes profound morphological changes, which are regulated by the cytoskeleton, including actin filaments, allowing it to deform to affect cell polarity, respond to mechanical cues and dictate migratory cell behaviours^46,47^. We further performed nuclear shape analyses in neurocrestoid-derived populations, and selected area, circularity, and solidity shape descriptors. We found that the nuclei of NCCs were smaller than those of other migratory cell types (average of 14.05% reduction in area), and they presented markedly decreased solidity and circularity values (N=3) (**Fig. S4B-C**).

We hypothesised that the characteristic nuclear shape of NCCs in our cultures might be conserved across species. To investigate this, we dissected mouse trunk neural crest explants and cultured them *ex vivo* for 48h as described in our previous publication^48^. A comparative analysis of nuclei from neurocrestoid-derived human NCCs and migratory NC from mouse explants (N=3) revealed similar nuclear morphology between the two species in terms of solidity and roundness. The primary difference observed was that human NCC nuclei were 26.92% larger than those of their mouse counterparts (**Fig. 4E-F**). Taken together, our results demonstrate that neurocrestoids can generate NCCs upon plating, including trunk NCCs, with distinct nuclear characteristics. This novel setup allows for human NC migration assays, which are directly comparable to mouse trunk NCC derived from explants.

### PDMS microgrooves allow directing, visualisation and filtering NCCs without altering their migratory capabilities

During migration, NCCs are subjected to a variety of cues which allow them to migrate through specific pathways while preventing invasion into restricted tissues. Amongst these, NCCs sense mechanical signals such as lateral confinement and substrate stiffening^49,50^, which are difficult to address with current human NC models. Moreover, in our cultures, NCCs frequently appeared in streams, with their trajectories constrained by adjacent SOX10- domains, reminiscent of the NC streams observed *in vivo* (**Fig. 4B** **and S4A**)^51^. However, the intermittent presence of restrictive SOX10-negative populations occasionally interfered with NCC migration, hindering the establishment of systematic migration analyses using plated neurocrestoids (**Fig. S4E**). To overcome these challenges, we incorporated PDMS micropatterned substrates into our neurocrestoid-plating system. Unlike most traditional microfluidic devices, PDMS substrates constitute open platforms that allow for imaging and serial manipulation of live long-term cell cultures, offering a simplified setup without the need for external pumps and flow control. Additionally, their relative ease of fabrication promotes broad adaptability and scalability, making them ideal for NC live migration assays^52^. To test their suitability for migration assays, we microfabricated PDMS microgrooves of different dimensions (width x height in µm): flat, 10x3, 10x10, 25x10 and 50x10. Following biofunctionalisation and Matrigel-coating to support cell culture compatibility, we plated organoids at day 8, allowing subsequent emigration of NCC (**Fig. 5A and C**).

**Fig. 5.**
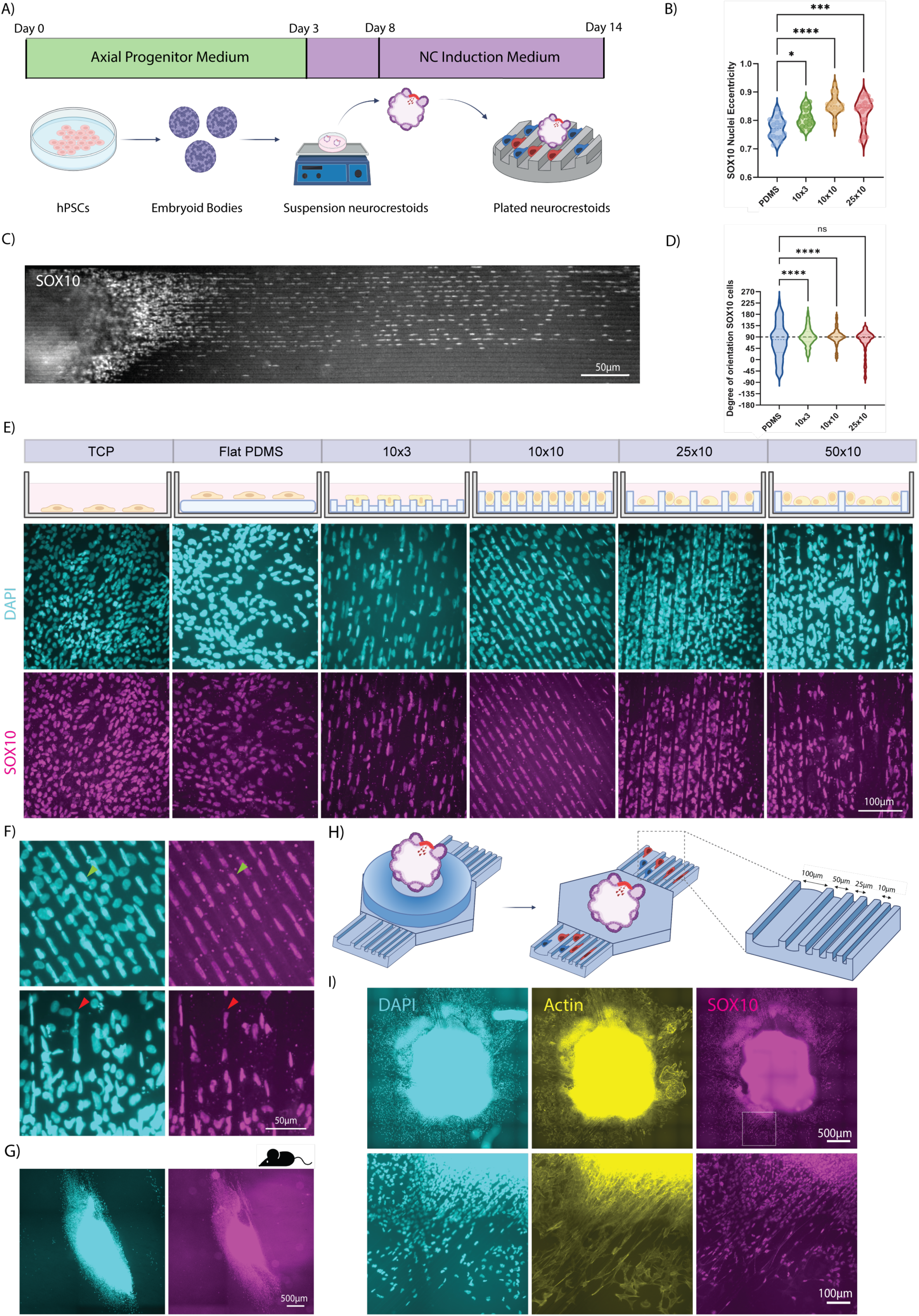
– PDMS microgrooves allow for live visualisation, guidance and selective migration of NCCs. A) Integration of neurocrestoids with PDMS micropatterned substrates allow for selective guidance and live visualisation of NC migratory populations. After 8 days in a rotary culture, differentiating neurocrestoids are plated in Matrigel-coated PDMS microgrooves for an additional 6 days of culturing. B) Violin plot showing the levels of eccentricity of SOX10-positive nuclei across different PDMS topographies. Datapoints represent the average eccentricity of all SOX10-positive nuclei within each field of view (n=5-250) in 15-17 technical replicates from 1 hiPSC line (Control 3) in 1 experimental block. Ordinary one-way ANOVA with Tukey’s multiple comparisons to test for significance between conditions. C) Representative image of the edge of a day 14 neurocrestoid plated on 10x10µm grooves. D) Violin plot depicting the degree of orientation of SOX10-positive nuclei of plated neurocrestoids in different PDMS topographies, where microgrooves have been set at 90°). Each violin plot represents a set of nuclei (n=5-250) in 15-17 technical replicates from 1 hiPSC line (Control 3) in 1 experimental block. Ordinary one-way ANOVA with Tukey’s multiple comparisons to test for significance between conditions. *P<0.5, **P<0.01, ***P<0.001, ****P<0.0001, ns= non-significant. E) Representative images of day 14 NCCs from plated neurocrestoids in 3x10, 10x10, 25x10 and 50x10µm PDMS grooves and flat PDMS and tissue culture plastics as controls. Samples are stained for SOX10 and nuclei (DAPI). F) Representative images of the area adjacent to the organoid core in neurocrestoids plated in 10x10 and 50x10μm PDMS microgrooves. Note that wider grooves can host SOX10-negative cells, resulting in restriction of SOX10-positive cell migration (red arrowheads). In narrower grooves, SOX10-negative cells do not fit, allowing for NCCs to migrate underneath along the grooves (green arrowheads). G) Representative fluorescence images after 48 hours of mouse trunk NC explants plated on 10x10µm grooves, accompanied by zoomed-in images of the areas delimited with a yellow dashed square. H) Schematic overview of the pipeline to plate neurocrestoids onto a PDMS multigroove star device. Organoids are positioned in the flat PDMS centre of the device using custom-made PDMS plating devices, which are removed once the organoids attach to the substrate. I) Representative images of a neurocrestoid plated in a multigroove PDMS device. The organoid is stained for SOX10, Actin (Phalloidin555) and nuclei (DAPI).

By day 14, we observed neurocrestoid-derived NCCs with the ability to interact and migrate along PDMS microgrooves. NCCs preferentially migrated through the microgrooves in all the different conditions, elongating their nuclei to fit inside the channels (**Fig. 5B-E**). Different-sized PDMS microgrooves provided NCCs with varying degrees of lateral confinement, with 10µm-wide grooves accommodating one single NCC at a time, while 25 and 50µm-wide grooves can host 2 to 5 SOX10-positive cells in parallel (**Fig. 5E**). We observed a significant reduction in the ability of SOX10-positive cells to migrate on flat PDMS versus control tissue culture plastics (approximately 47%), potentially suggesting a preference of NCCs for stiffer substrates. Microgroove topographies were partially able to reverse this effect, with wider grooves showing a modestly higher migration rate (average 325µm further than narrow grooves, N=3) (**Fig. S5A**). Importantly, we noticed that SOX10-negative populations, which were previously noted to restrict NCC migration, were present inside the wider topographies but did not fit into the 10x10µm grooves (**Fig. 5F**). SOX10-positive cells represented 70.67% of the cells that migrated through the narrower channels (**Fig. S5B**). Moreover, we trialled our PDMS microgrooves on trunk NC mouse explants and observed successful migration along grooves of different topographies after 48 hours, opening the door for direct cross-species NCC migration comparisons (**Fig. 5G** and **S5C**). Thus, we concluded that the narrow, 10x10µm, groove design provides a system that allows for imaging of selective guided NCC migration irrespective of other cell types present and constitutes the standard for future analyses.

One of our original goals was to overcome the potential variability between organoids and customise a system to assess multiple migratory conditions in parallel. Therefore, we designed a new bioengineered device where NCCs derived from a single organoid could migrate into channels of widths of 10, 25, 50, and 100μm. Organoids were seeded in the hexagonal surface using custom-made PDMS stencils, and delaminating NCCs migrated radially into the different topographies (**Fig. 5H-I** **and S5D**). Analysis of the total distance travelled revealed that groove width was not a determining factor for individual NCC migratory efficiency, as demonstrated by similar migration rates by SOX10-positive cells across different topographies (N=3) (**Fig. S5E**).

Overall, by integrating micropatterned PDMS substrates with our neurocrestoid plating strategy, we generated a novel hPSC-derived migration-on-chip platform, which allows for selective direction and live imaging of NC migratory populations derived from a three-dimensional organoid. Importantly, this platform is highly amenable and can be easily customised in terms of topography and mechanical properties for future human NC migration studies, including the effect of mechanical stress and durotaxis. We have demonstrated that cell-intrinsic mechanisms override lateral confinement but not substrate stiffness in regulating hPSC-derived NC migration. Finally, we demonstrated the ability of narrow grooves (10x10μm) to direct NCC migration and isolate them from other restrictive SOX10-negative cell types. Altogether, our neurocrestoids recapitulate key developmental stages of NC development in a complex 3D environment, which we were able to validate using ex-vivo mouse embryo. Neurocrestoids can be integrated with micropatterned PDMS substrates to generate a novel, powerful migration-on-chip platform to systematically study human NCC migration from hPSC-derived organoids.

## Discussion

Human NC development has long been challenging to study due to the scarcity of early embryonic samples and the lack of appropriate models. In recent years, many studies have developed protocols to derive NCCs from hPSCs in 2D monolayer cultures^29,32,34,53,54^. However, these lack the cellular and structural complexity necessary to investigate key aspects of NC development. A very attractive approach to modelling human development is the use of hPSC-derived 3D organoids, which mimic key aspects of development in a multicellular environment, showcasing complex architectures. However, the availability of organoid protocols in the field of NC is limited. An established pipeline in the field relies on spontaneous aggregation of hPSCs to generate neurospheres capable of producing anterior NCCs upon plating^17,18^. Similarly, Kirino and colleagues established a protocol to generate sympathetic neurons from neural spheres^55^. However, none demonstrated the generation of NCCs in 3D. In recent years, several new studies have described the generation of anterior NC organoids, but these are restricted to the cranial compartment^21,22^.

In this study, we have described a novel bioengineered protocol to generate neuromesodermal organoids capable of producing an array of NCCs along the anteroposterior axis, including trunk NCCs. These neurocrestoids express hallmarks of NC development, including NMP generation, NPB establishment, NC identity acquisition, embryonic patterning and EMT. Neurocrestoids possess tissue architecture reminiscent of *in vivo* NC, displaying a hierarchical tissue-specific progression of SOX9 in the neuroepithelium followed by SOX10 in delaminating NC. Our organoids generate abundant migratory populations of NCCs, which can then be plated on PDMS micropatterned microchannels suited for selective isolation, guidance and live visualisation in migration assays. Importantly, our results demonstrate the generation of NCCs along the anteroposterior axis, including cranial, vagal and trunk NCCs, complementing previous studies that focused solely on generating the cranial crest and responding to a pressing need for the generation of 3D *in vitro* models for posterior NC development.

The custom-made PDMS microwells shown here are similar to those in our recent publication^26^, which allow for high-throughput generation of EBs of a regular size, helping to standardise organoid cultures and reduce variability. Importantly, these microwells are also customisable; for example, we have continuously updated our microgroove design to adapt it to differently-sized well plate formats (24 and 48 wells) according to our experimental needs. In contrast to traditional EB aggregation methods, which are generally low-throughput and expensive commercial systems limited by predefined well dimensions, plate formats, and geometries, our PDMS microwell approach offers a cost-effective and highly customizable alternative, and is readily amendable to further scaling. However, we observed another small population (12.4%) of EBs with diameter σ > 1%, which results from the fusion of individual aggregates. To circumvent this problem, further redesign of our microwells could be performed to increase well depth and avoid EBs escaping the pockets and fusing with their neighbours.

Our data suggest that neurocrestoids contain most of the major cell types present at the time of neural crest development, including neural ectoderm, NC, cranial placodes and paraxial mesoderm, except non-neural ectodermal (data not shown), likely due to the early addition of FGF2^56^ and sustained CHIR-mediated Wnt activation^57^. *In vivo*, juxtaposition of such tissues shapes NC development by establishing gradients of signalling molecules, which activate the Wnt, BMP, FGF, and Notch pathways, and ultimately determine NC fate specification^4,37^. Consistent with this, our data show timely upregulation of genes encoding specific components of these pathways, many of which are secreted *in vivo* by surrounding tissues and act as essential regulators of NC development. Amongst these, we demonstrated that modulation of Wnt activation during the first 3 days of the protocol was paramount for NC production, organoid cell composition and architecture, confirming previous findings in 2D adherent cultures^29,34,36^. We believe the neurocrestoid protocol to be robust since it has been validated amongst 4 distinct hPSC lines across two independent laboratory sites (The Francis Crick Institute and King’s College London). Taken together, these experiments highlight the importance of generating *in vitro* models that incorporate the major relevant tissues, allowing the study of NC cells not in isolation, but within the context of a rapidly developing and dynamically changing environment. Further characterisation of non–cell-autonomous mechanisms driving NC developmental events in the human embryo using neurocrestoids represents an important avenue for future research.

One caveat of our system is a modest temporal and structural variation in neurocrestoids across batches and cell lines (i.e. morphological differences in **Fig. 1A** versus **S1E** or variations across replicates in **Fig. 2**), a common challenge amongst organoid protocols^58,59^. This inherent variability might partly result from fluctuations in the proportions of non-NC populations (mesoderm, neural ectoderm) arising from early stochastic events during the differentiation processes. Shifts in these cellular ratios can alter endogenous signalling dynamics and thereby can influence lineage decisions. This is particularly critical for the NC, which arises from the neural–non-neural ectoderm boundary and depends on tightly regulated intermediate levels of signalling molecules, mostly Wnts and BMPs, with even subtle modifications redirecting fate towards neural, non-neural, or placodal fates^57,60^. This constitutes an additional layer of complexity, underscoring the need for the development of strategies that maximise experimental reproducibility, such as the use of bioengineering techniques.

Due to the multicellular nature of neurocrestoids, systematically evaluating migratory behaviour of NCC can be challenging in a conventional culture, and the presence of “restrictive” cell types (**Fig. S4**) can further complicate this. For these reasons, we integrated our 3D system micropatterned PDMS channels to selectively guide NC migration and improve current human NCC migration assays, obtaining a “migration-on-chip” specific platform for human NCCs. By plating neurocrestoids at day 8 in narrow (10x10μm) microgrooves, we can generate, visualise, and direct NCC migration, irrespective of other restrictive cell types. This platform is easily customisable and opens the door for systematic human 3D-generated NCCs in health and disease. We anticipate that in future studies, incorporating genetic reporters such as a SOX10 live fluorescent reporter hPSC line will position our migration-on-chip platform to be a powerful tool to study human NCC dynamics both in normal and pathological contexts.

In conclusion, we report a novel platform, which combines the use of hPSCs, 3D organoid cultures and bioengineering to study human NC development. These organoids are capable of producing an array of NC subtypes, including trunk NCCs in 3D associated with physiologically relevant epithelial structures and EMT behaviours. Neurocrestoids recapitulate hallmarks of NC development and contain most of the major relevant ectodermal and mesodermal subtypes required for NC induction. By integrating the use of PDMS microgrooves, we have created a migration-on-chip platform enabling the production, direction and live visualisation of NCC migration irrespective of other cell types. In light of our limited understanding of human NC development, the importance of the complex 3D environment in which NCCs develop and the lack of appropriate models, we anticipate that neurocrestoids will represent a ground-breaking system for studying NC development along the anteroposterior axis in health and disease.

## Materials and Methods

### Stereolithography 3D-printing

PDMS microwells (400x400x250µm) for EB formation, plating devices (600µm radius at the base) and slide spacers (500µm thickness) for organoid mounting (1000x1000x1000µm at the base) were generated from 3D-printed resin moulds as per our latest publication^26^. Microwells were redesigned using Fusion360 (Autodesk) by increasing the well depth from 125µm to 250µm to avoid contiguous EB fusion, while avoiding the formation of air bubbles, which persist even after high-speed centrifugation present in deeper well designs. The total surface covered by the mould features was also increased to 3300µm in length to maximise the number of microwell devices obtained per cast, while ensuring fitting within the area of a microscope slide at the time of clipping.

Mould designs were sliced using Chitubox (3D Slicer Software), 3D printed and post-processed as previously described^26^. Printer model, settings and resins used can be found in Supplementary Table 1. Mould dimensions were measured using a 3D Optical profiler (Sensofar S Neox) and resulted in 40-50µm shallower wells when compared to the CAD design, likely due to the inherent limitations in the printer resolution.

PDMS elastomer was mixed in a 10:1 ratio with polymerising agent (SYLGARD 184, Dow) and degassed for 1 hour on a vacuum desiccator (Sigma, Z119016) to remove air bubbles. To ensure complete removal of any resin excess, we performed a first cast by pouring PDMS onto the newly-printed moulds and baked for 1 to 5 hours at 75°C, which was always discarded. Subsequent casts were performed by pouring the PDMS mixture into the moulds, clipping them using two microscope glass slides, and baking for 15 minutes at 75°C. Once the PDMS was cured, devices were demoulded and UV-treated for 15 minutes prior to any cell culture applications.

### Photolithography

Micropatterned silicon wafers used to generate PDMS microgrooves were fabricated by photolithography as previously described^52^. First, designs were created using CleWin5 (Layout Editor Software), and a custom emulsion photomask (0.18mm) was generated by JD PhotoData. Test grade N(phos) wafers (Pi-Kem) were pre-treated with Oxygen plasma (10 minutes, 7sccm, 100% power) in a plasma cleaner (Henniker Plasma HPT-100) and baked for 20 minutes at 200. The epoxy-based negative photoresist, SU8-2007 (KAYAKU Advanced Materials), was layered on the silicon wafer using a spin coater (POLOS200 Advanced, SPS) at 1,500 rpm. Layered silicon wafers were left to rest for 5 minutes, softbaked for 1 minute at 65°C and for 3 minutes at 95°C. Following pre-treatment, we placed SU-8-layered silicon wafers in the mask aligner (UV KUB 3, kloe-france) and exposed them to UV light through the photomasks for 8 seconds at 50% power (130mj/cm^2^). In order to fully fixate microfeatures, we post baked the newly micropatterned silicon wafers for 1, 4 and 1 minute at 65, 95 and 65°C, respectively. Finally, the remaining SU-8 was washed off in a shaker for 4min in PG-MEA (Sigma, 484431), washed with isopropanol and then silicon wafers were hard-baked for 20 minutes at 200°C.

To facilitate demoulding and ensure that microfeatures would remain after PDMS casting, micropatterned silicon wafers were silanized. First, Oxygen plasma-treated (5 minute, 7 sccm, 100% power). Wafers were subsequently placed on a vacuum desiccator along with 200μl of Trichlorosilane (Sigma, 448931) for 1 hour. Finally, the excess of silane was washed off using Acetone (Sigma, 179124).

### Soft-lithography

PDMS micro and multigrooves were generated as previously described^52^. PDMS monomer was mixed at a weight ratio of 10:1 to the curing solution, desiccated, spun at 300rpm on the silicon wafer using a spin coater (Polos) and baked for 5 to 10 minutes at 100°C. To ensure biocompatibility, substrates were oxygen plasma treated (30s, 100%, 7sccm), coated with Poly-D-Lysine 0.01% and Matrigel overnight unless otherwise stated prior to organoid plating.

### Human induced pluripotent stem cell culture

HiPSC lines were obtained either from commercially available sources: Control 3 (ThermoFisher Scientific, A18945), KOLF2.1J (Jackson Laboratory, JIPSC1000), or kindly donated: KOLF2C1 (Maximiliano Gutierrez, The Francis Crick Institute). The HESC line used was H9 (WiCell, https://hpscreg.eu/cell-line/WAe009-A-2J).LHPSCs were cultured in Matrigel-coated plates (Corning, 356234) and fed with Essential 8 Flex medium (ThermoFisher Scientific, A2858501) at 5% CO2 37°C. Once confluent, hPSCs were passaged using a cell dissociation buffer (ThermoFisher Scientific, 13151014) for 3 minutes at 37°C. HPSCs were regularly tested for mycoplasma and genetic abnormalities.

### Neurocrestoid generation

Microwell plates were sterilised using a UV lamp for 15 minutes and centrifuged at 4000rcf for 5 minutes in day 0 differentiation medium (D0DF, Supplementary Table 1) supplemented with 10μM Y-27632-dihydrochloride to avoid bubble formation inside the wells. CHIR99021 concentration ranged between 0.5μM (KOLF2C1) to 1μM (CTL3 and KOLF2.1J) and had to be optimised for each hPSC line and CHIR batch. When hPSC colonies reached a diameter of approximately 800µm, cells were singularized using Accutase for 10 minutes at 37°C and seeded at 700,000 cells per 48-well plate stamp of our custom-made 400x400x250µm PDMS microwells. Once seeded, plates were centrifuged again at 100rcf for 3 minutes to capture the cells inside the wells.

After 18 hours, we withdrew Y-27632. After 24 hours, EBs were collected and transferred to an orbital shaker at 60rpm in 60mm-Petri dishes until day 3. On this day, medium was changed to day 3 differentiation medium (D3DF, Supplementary Table 2) plus 0.5-2% Matrigel, depending on the Matrigel batch and organoids were fed every other day until plated or until the endpoint of each experiment. Brightfield images of the differentiation process were acquired using an EVOS M5000.

Neurocrestoids were plated at different timepoints during differentiation in Matrigel-coated 96-well plates with or in 48-well plates containing PDMS micropatterned substrates in D3DF with 10μM Y-27632. The next day, Y-27632 was withdrawn, and neurocrestoids were fed every other day.

### Genetically-modified mouse models

Animal work was approved by King’s College London Animal Welfare and Ethical Review Body (AWERB) and performed at King’s College London in accordance with the UK Home Office Project Licenses PP1672528 (TS) and PP3807977 (TS). Tg(Wnt1::cre)11Rth mice (MGI allele ID 2386570) as previously described in^61^ were crossed with the reporter line Rosa26Rmtmg (GT(Rosa)R26SorTm4(ACTB-tdTomato-EGFP)Luo) (MGI allele ID 3716464)^62^. All mouse lines are maintained on a Crl:CD1(ICR) outbred background. After mating, the observation of a vaginal plug was considered embryonic day 0.5 (E0.5)

### Neural crest explant cultures

At E9.5 (24 to 27 somites), pregnant females were culled and the uterus was removed and placed on ice-cold PBS (Sigma-Aldrich, D8537) in a 10cm dish. The mesometrium of the uterus was cut, and the muscle layer was removed to separate the individual decidua. The neural plate border (NPB) adjacent to somites 4-6 from the tail end was dissected out. The NPB was divided down the anteroposterior axis so that each side of the neural plate border (explant) could be plated individually. Explants were transferred into a 35 mm coverslip glass-bottomed dish (Ibidi, 82426) and incubated at 37°C and 5% CO_2_. Each well was pre-coated with 1 μg/mL fibronectin (Sigma-Aldrich, F1141) and the explants were cultured in neural crest media (DMEM-high glucose (Sigma-Aldrich, D5671), 15% embryonic stem cell-grade fetal bovine serum (Thermofisher, 16141079), 0.1mM minimum essential medium nonessential amino acids (Gibco, 11140), 1mM sodium pyruvate (Gibco, 11530396), 55μM β-Mercaptoethanol (Gibco, 31350), 100 units/mL penicillin, 100 units/mL streptomycin (Sigma-Aldrich, A5955) and 2mM L-Glutamine (Scientific Laboratory Supplies, G7513), conditioned by growth-inhibited STO feeder cells (ATCC) and supplemented with 25 ng/μL basic-FGF (R&D System, 233-FB-025/CF) and 1:1000 LIF (Sigma-Aldrich, ESG1106).

### Whole organoid staining, clearing and analysis

Neurocrestoids were fixed with 4% PFA (Boster, AR1068) for 1 hour and washed three times with PBS for 30 minutes. Permeabilisation and protein blocking were performed in one step using a solution containing 5% (Sigma, G9023) and 1% Triton-X (Sigma, T8787) in PBS and incubated for 5 hours. Primary antibodies were prepared in an antibody solution consisting of 0.5% Triton-X and 5% Goat serum and applied overnight at 4°C. Primary antibodies were washed three times with PBS for 30 minutes. Secondary antibodies were prepared in the same antibody solution and incubated for 2 hours. Finally, the excess was washed three times with PBS. All antibodies and their concentration can be found in Supplementary Table 3. Cell nuclei were stained using 1ng/mL of DAPI (Thermofisher Scientific, D1306). All steps were performed in a platform shaker at 30rpm at room temperature unless otherwise stated.

Neurocrestoids were placed and cleared between microscope slides and coverslips using custom-designed PDMS spacers consisting of a 500µm-thick layer of PDMS with 3.25mm diameter circular overtures. Clearing was performed using a single-step clearing agent (Rapiclear 1.47, Sunjin Lab, RC147001) for 24h at room temperature. Neurocrestoids were imaged using an Olympus CSU-W1 SORA spinning-disk system. 3D rendering images and optical slices were visualised using Imaris software.

### Immunostaining of 2D cultures and analysis

Plated neurocrestoids and all other 2D cultures were fixed using 4% PFA for 15 minutes and washed three times with PBS. Cells were permeabilised with 0.4% Triton-X for 10 minutes and incubated with blocking solution containing 3% goat serum for 30 minutes. Primary antibodies were prepared in blocking solution and incubated overnight at 4°C. Once washed three times with PBS, secondaries were applied in blocking solution for 1 hour and washed again prior to mounting. A list of all antibodies can be found in Supplementary Table 3. Actin was stained using ActinGreen488 ReadyProbes (Thermofisher, R37110) for 5 minutes at RT or along with the secondary antibodies and washed 3 times to remove the excess of dye. Fluorescence imaging was performed on a Nikon Ti2 Eclipse system.

### *In situ* hybridisation chain reaction and confocal imaging

*In situ* hybridisation chain reaction (HCR^TM^) was carried out as standard protocol for mouse embryos by Molecular Instruments, as per^63^. Each custom-designed probe was added to the final concentration of 4nM in hybridisation buffer and incubated at 37°C. Hairpins were separately heated (30pmol each) at 95 C for 90 seconds, cooled in a dark drawer for 30 min at room temperature, and then combined in 500μl of amplification buffer. Embryos are incubated with hairpin solution protected from light, overnight at room temperature. Samples are embedded in 1% low-melting-point agarose (Sigma, A4018), cleared with Ce3D^TM^ (Biolegend, 427704) and kept protected from light and taken to be imaged. Images were taken with an inverted Nikon A1R confocal microscope.

### Visualisation and analysis of adherent cultures

Fluorescence images of adherent cultures, including plated organoids and mouse trunk NC explants, were visualised and analysed using Fiji/ImageJ software.

Percentage of SOX9, SOX10 or HOXC9 positive area analysis was performed using the “polygon selection” tool over the DAPI channel (nuclei), exclusively including the total area covered by the plated organoid and its migratory cells. We display the percentage of the area that cells positive for the protein of interest cover over the total DAPI area.

Morphology analysis comparing NC versus non-NC populations and human and mouse NC nuclei was performed manually on images blinded using the “blind analysis tools”. We randomly selected and delineated using the polygon selection tool 6 to 10 nuclei, and their shape parameters were measured without consideration of their SOX10 status. In the case of the NC versus non-NC nuclei morphology comparisons, SOX10 positivity was always checked after measurement and cells were classified accordingly.

NCC occupation was calculated using the “Cell Counter” ImageJ plugin and the “polygon selection” tool. We randomly delineated the space corresponding to a single microgroove on the brightfield channel and counted the number of nuclei present in the DAPI and SOX10 channels. Results are presented as the number of NCCs out of the total cells that occupied each PDMS microgroove.

To measure the efficiency of migration of NCCs within plated neurocrestoids and mouse trunk explants, we selected the five most distant NCCs from the organoid edge from populations containing at least 4 cells for each organoid (single NCCs were excluded), using the “segmented line” tool. For analyses showing distance ratios, the average distance was calculated for the control condition, and each value was normalised to this parameter.

To quantify eccentricity and orientation of nuclei within PDMS microgrooves, we designed a semi-automated pipeline integrating Ilastik, ImageJ/Fiji and CellProfiler. We trained Ilastik for image segmentation on a random subset of images for each channel (approximately 10%). Thus, pixel classification was user-independent and accurately classified by the software. Once the training phase was completed, all images were batch processed to generate binary segmentation images automatically. This process was repeated for each channel (DAPI, SOX10) for an accurate pixel segmentation. Binary segmentation images were transformed into binary masks using a custom-made ImageJ macro, and these were fed into a custom-made CellProfiler pipeline. This pipeline was designed to identify objects within the binary masks for each channel, label and quantify the number and morphology parameters of such objects. NC nuclei were obtained by superimposing the SOX10 and DAPI binary masks. Non-NCC were objects only present in the DAPI channel, that fit into the nuclei size criteria.

For SOX10-positive nuclei orientation analysis, a rotation correction was applied to account for variations in the alignment of PDMS grooves across wells. Grooves were rotated to 90° (vertical alignment), and the corresponding rotation angle was subtracted from all the individual values for that field of view. Finally, a directionality analysis was run on ImageJ.

### Gene expression analysis

Total RNA was extracted using the RNAeasy Mini Kit (Qiagen, 74104) following the manufacturer’s instructions. Subsequent retrotranscription to obtain cDNA was performed using the High-Capacity RNA-to-cDNA kit (Thermofisher Scientific, 4387406). Real-time quantitative PCR reaction was run on a 7500 Real-Time PCR system (Applied Biosciences) using Fast SYBR Green Master Mix (ThermoFisher Scientific, 4385612). Relative gene expression analysis was obtained by ΔΔCt method normalized to the housekeeping gene GAPDH, and using day 0 hiPSCs as a control. Primer sequences used can be found in Supplementary Table 4.

### Statistical analysis

As a first step, normality of the samples was assessed using a Shapiro-Wilk test. Subsequently, a corresponding parametric or non-parametric test was chosen. For 2 non-parametric groups, the Mann-Whitney test was applied. For 3 or more parametric samples, a one-way ANOVA accompanied by Tukey test for multiple comparisons was used. For non-parametric analysis, a Kruskal-Wallis and Dunn’s for multiple comparisons were performed. Graphs were obtained using custom R files or Graphpad Prism. Statistical significance has been represented as *,**,***,**** for p-values of <0.05, <0.01, <0.001, <0.0001, respectively.

### Library Preparation for Bulk RNA-sequencing

Neurocrestoids were generated in three technical replicates from the Control 3 hiSPC line as previously described, each deriving from a separate well of the stem cell culture. Longitudinal samples were obtained from 20 differentiating organoids per replicate at days 0, 3, 6 and 8. Additionally, NCCs were sorted from day 8 neurocrestoids using p75 magnetic beads as per the manufacturer’s instructions (Miltenyi Biotec, 130-099-023).

Samples were lysed in 1ml of TRIzol Reagent (Thermofisher Scientific, 15596026) and stored at -80°C until further processing. Total RNA for bulk RNA sequencing analysis was extracted using TRIzol Reagent as per the manufacturer’s instructions.

A total of 15 samples containing 2μg each of RNA were sent to Novogene for RNA sample quality control, mRNA library preparation (non-strand specific with polyA enrichment), RNA sequencing using Illumina NovaSeq X Plus Series (PE150, 50M reads) and data quality control.

### Bulk RNA-seq data processing

Primary data analysis was performed using the nf-core/rnaseq pipeline (version 3.19.0) [https://github.com/nf-core/rnaseq] with default parameters written in the Nextflow domain-specific language (version 24.10.2, build 5932), and using the Singularity configuration. The fastq files were aligned to the human reference genome (GRCh38.dna. primary_assembly) and annotated using the GRCh38-114 release GTF from Ensembl. The gene-scaled counts matrix output from STAR_salmon was further filtered by removing genes that had less than 10 counts found across less than 3 samples.

### Differential expression analysis

Differential expression analysis was run using DESeq2 (version 1.48.1) in R (version 4.5.1) using the standard workflow. In the creation of the DDS object, the counts were rounded to the nearest count, and the condition in this case was the day of harvesting (D0, D3, D8 or P75 sort). Principal component analysis (PCA) plots were plotted following variance stabilisation, highlighting the cells’ condition. Differential expression analysis was conducted by fitting negative binomial generalised linear models and assessing for significantly changed genes using Wald statistics. Significance was defined by an adjusted p-value of less than 0.05 and an absolute log2 fold-change of at least 1. Scaling was achieved using the basic R scaling function on the DESeq2 normalised gene counts. All filtered significantly differentially expressed genes were compiled into an Excel file using the openxlsx package. Heatmaps were generated using the ComplexHeatmap package (version 2.24.1). All remaining plots were generated using ggplot2 (Version 3.5.2).

## Supplementary Figures

**Fig. S1.** - Development of a bioengineered pipeline for the generation of neuroepithelial organoids capable of producing NCCs in 3D and upon plating using custom-made bioengineered microwells. A) Schematic of the pipeline followed to generate custom-made PDMS microwells as described (Hagemann et al., 2024). B) Representative brightfield images showcasing previous issues in spheroid formation in the previously published PDMS microwell designs: in 400x400x125 microwells, spheroids escape the pockets and fuse (yellow arrowheads), while in the 800x800x400 design, we observe significant air bubble formation (pink arrowhead). C) Optical profile measurement of a microwell resin mould demonstrating the real dimensions of PDMS microwells to be 210µm in depth. D) Representative optical slices of whole-mount immunostained neurocrestoids derived from three additional hPSC lines: H9 ESCs and KOLF2.1J hiPSCs at Guy’s Hospital and KOLF2C1 hiPSCs at The Francis Crick Institute, highlighting the robustness of our protocol. E) Representative Z-stacks of CTL3-derived organoids treated with low (1-1.5µM), medium (2-2.5µM) or high (3-3.5µM) CHIR dosages for the first 3 days of the differentiation protocol. White dashed lines delimit the epithelial structures within organoids. F) Representative images of day 14 plated neurocrestoids from the low, medium and high CHIR conditions and immunolabeled for SOX10. G) Quantitative gene expression analysis of SOX2, CDX2, TBXT and SOX10 at day 8 neurocrestoids treated with increasing CHIR concentrations (1-3.5µM). Relative mRNA expression levels were quantified by qPCR and performed in technical duplicates. Data normalised to GAPDH and presented as a fold-change versus hiPSCs (day 0), mean ± S.D. H) Bar chart showing percentage of SOX10 positive cells amongst the total nuclei (DAPI) in plated neurocrestoids treated with low (1.5µM), medium (2.5µM) or high (3.5µM) CHIR concentrations (days 0 to 3), mean ± S.D. Datapoints represent single organoids (n=26-30) in 6-11 technical replicates from 2 hiPSC lines (Control 3 and KOLF2.1J) in 3 experimental blocks. Non-parametric Kruskal-Wallis test with Dunn’s multiple comparisons to test for significance between days, ns = non-significant. ****P<0.0001, ***P<0.001.

**Fig. S2.** – Neurocrestoids can recapitulate the key signals involved in NC development and suggest transcriptional signatures of posterior NC derivatives. A) Principal component analysis plot of the transcriptomics dataset showing day 0, 3, 6, 8 and p75-sorted samples clustering by condition. B) Dot plots showing the normalised counts for p75NTR, SOX10 and PAX6, demonstrating that magnetic bead p75 sorting allows isolation of NCCs from 3D suspension neurocrestoids. C) Bar charts depicting the number of differentially upregulated and downregulated genes in each condition compared to the previous timepoints or to day 0. D) Heatmap of differentially expressed genes (significant padj<0.05 |log2FC|>1; D8 vs D0) relating to sympathetic neurons (SYMPN), Schwann cells and Schwann cell precursors (SC/SCP), melanocytes (MLANO) and sensory neurons (SENSO) in the day 8 versus day 0 conditions.

**Fig. S3.** – Neurocrestoids show transcriptional signatures hallmark of embryonic organ development and anteroposterior tissue patterning. A) Panel showing the MA plot, a heatmap of the most differentially expressed genes and GO biological process enrichment analysis for comparison between P75-sorted and day 0 conditions. B) Corresponding analyses (MA plot, heatmap of the most differentially expressed genes, and GO Biological Process enrichment) for the comparison between day 8 and day 0 conditions. C) Corresponding analyses (MA plot, heatmap of the most differentially expressed genes, and Gene Ontology (GO) Biological Process enrichment) for the comparison between P75-sorted and day 8 conditions. MA plots show the log10 of the base mean expression plotted against the log2 Fold Change in gene expression. Each dot represents a gene, with significantly differentially expressed genes highlighted in red. The blue dotted lines showcase a differential gene expression of |log2FC|>1, which is the threshold for transcripts to be taken into consideration for downstream RNA-seq analysis in the comparisons between each pair of conditions. Heatmaps show the top 50 most differentially expressed genes (up- and downregulated) for each comparison. GO Biological Process (BP) enrichment analysis for each set of condition comparisons. Red indicates a GO term related to genes upregulated in the first condition, and blue indicates downregulated.

**Fig. S4.** – Neurocrestoid-derived NCCs show distinct nuclear and cellular characteristics versus SOX10-negative populations, which restrict their migration. A) Representative fluorescence images at day 14 of plated neurocrestoids derived from KOLF2C1 and KOLF2.1J hiPSC lines stained for SOX10 and nuclei (DAPI), accompanied by zoomed-in images of the areas delimited with a yellow dashed square. B) Representative images of SOX10 negative and positive populations, which served for subsequent nuclei shape analyses. The last column displays examples of nuclei with a higher (round) versus lower (beam) circularity and solidity indexes. C) Bar charts indicating the area, circularity and solidity values for SOX10 positive versus negative cells, mean ± S.D. Non-parametric Mann-Whitney test to test for significance between conditions. *P<0.5, ****P<0.0001. Datapoints represent average values for all the SOX10-positive or negative cells in a given field of view. Total number of points was 38-51 per condition, corresponding to 5 fields of view per organoid, 3-5 organoids per cell line from 3 hiPSC lines (Control 3, KOLFC21 and KOLF2.1J) in 3 experimental blocks. D) Representative immunofluorescence images of the migratory cells around a plated neurocrestoid at day 14, showcasing both SOX10-positive and negative populations. Cells are stained for Actin (ActinGreen488), SOX10 and nuclei (DAPI). White arrowheads indicate the distinct Actin localisation patterns between both populations. E) Representative immunofluorescence image of a neurocrestoid plated in a standard tissue culture plastic plate depicting SOX10-negative areas restricting NCC migration (white arrowheads).

**Fig. S5.** – Narrow 10x10µm grooves selectively guide NCC migration but do not alter their migratory capabilities. A) Bar chart depicting the distance migrated by SOX10-positive cells from the organoid core for each topography and TCP as a control, mean ± S.D. Each point represents the average distance by SOX10-positive cells in each organoid (n = 33-25), calculated using 3-5 cells per organoid in 7-14 technical replicates in 2 hiPSC lines (Control 3 and KOLF2C1) in 3 experimental blocks. Ordinary one-way ANOVA with Tukey’s multiple comparisons to test for significance between conditions. **P<0.01,****P<0.0001. B) Pie chart showing the average percentage of SOX10-positive versus negative cells that localise along each single 10x10µm groove. C) Representative fluorescence images after 48 hours of mouse trunk NC explants plated on 10x3, 10x10 and 50x10µm grooves and flat PDMS, accompanied by zoomed-in images of the areas delimited with a yellow dashed square. D) Zoomed-in images of the organoid plated on star multigroove device on Fig. 5H stained for SOX10, Actin and DAPI. E) Bar chart showing the distance travelled by SOX10-positive cells in multigroove-plated neurocrestoids at day 14 through the different groove widths, mean ± S.D. Datapoints represent average distance migrated by SOX10-positive cells for each organoid (calculated using 3-5 cells per organoid) (n = 33-25) in 7-14 technical replicates in 2 hiPSC lines (Control 3 and KOLF2C1) in 3 experimental blocks. Non-parametric Kruskal-Wallis test with Dunn’s multiple comparisons to test for significance between widths, ns: non-significant. F) Brightfield image of PDMS multigroove substrate showcasing the differently-sized microgrooves side by side.

## Supplementary Tables

**Table 1:**
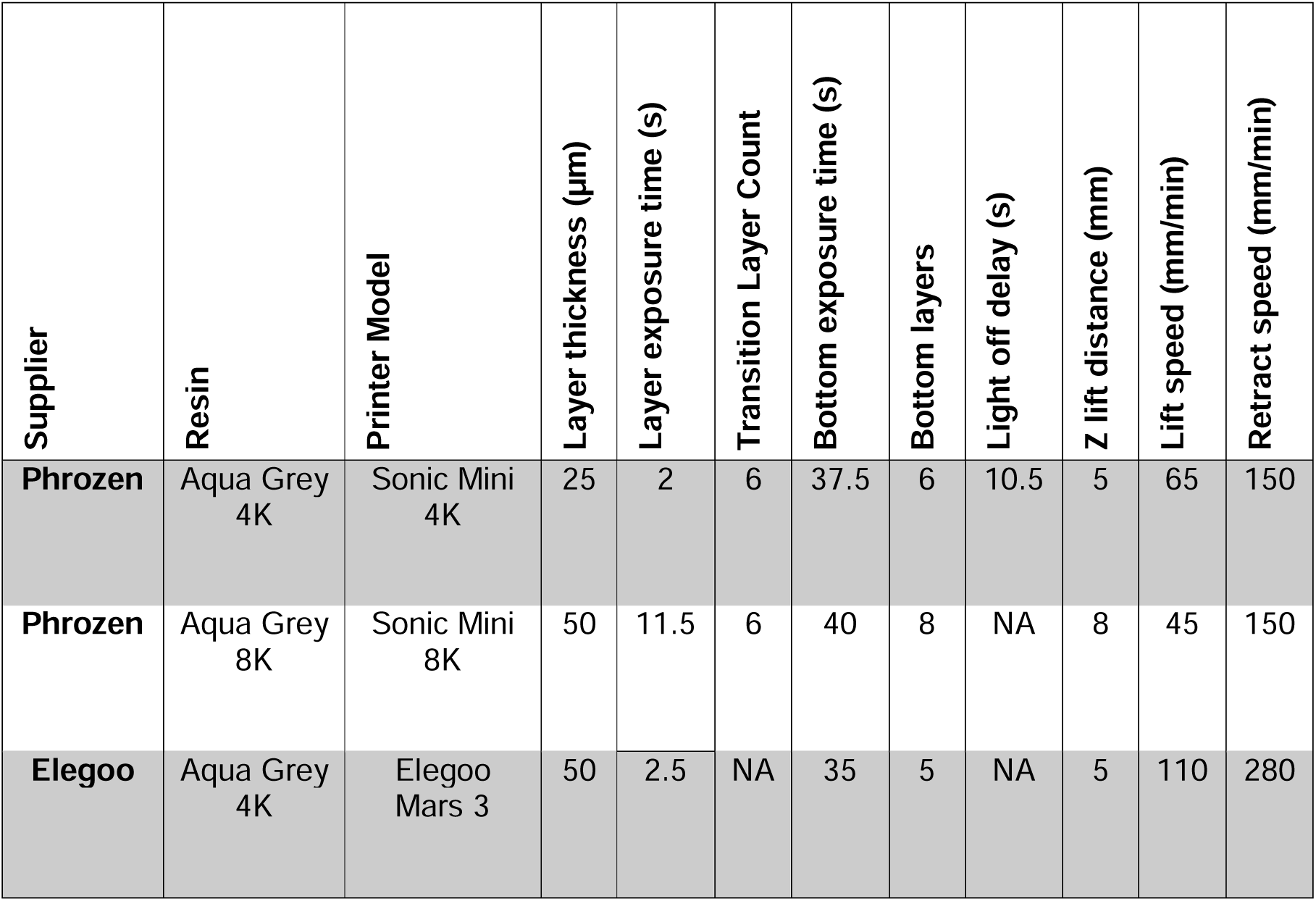
Printer settings, model and resin used to print PDMS microwells, plating devices and slide spacers.

**Table 2:** Day 0 differentiation medium recipe.

| **Reagent** | **Final Concentration** | **Brand** | **Catalogue Number** |
| --- | --- | --- | --- |
| **Advanced DMEM/F12** | 1:1 | ThermoFisher Scientific | 12634010 |
| **Neurobasal** | 1:1 | ThermoFisher Scientific | 21103049 |
| **B27** | 1X | ThermoFisher Scientific | 17504044 |
| **N2** | 1X | ThermoFisher Scientific | 17502048 |
| **Glutamax** | 1X | ThermoFisher Scientific | 35050061 |
| **Penicillin-Streptomycin** | 1X | ThermoFisher Scientific | 15070063 |
| **100x MEM Non-Essential Amino Acids** | 1X | ThermoFisher Scientific | 11140050 |
| **CHIR99021** | 0.5µM - 9µM | Tocris | 4423 |
| **FGF2** | 20 µg/mL | ThermoFisher Scientific | 13256-029 |

**Table 3:** Day 3 differentiation medium recipe.

| Reagent | Final Concentration | Brand | Catalogue Number |
| --- | --- | --- | --- |
| Advanced DMEM/F12 |  | ThermoFisher Scientific | 12634010 |
| N2 | 1X | ThermoFisher Scientific | 17502048 |
| Glutamax | 1X | ThermoFisher Scientific | 35050061 |
| Penicillin-Streptomycin | 1X | ThermoFisher Scientific | 15070063 |
| 100x MEM Non-Essential Amino Acids | 1X | ThermoFisher Scientific | 11140050 |
| CHIR99021 | 1μM | Tocris | 4423 |
| SB431542 | 1μM | Tocris | 1614 |
| Recombinant BMP4 | 15ng/ml | ThermoFisher Scientific | PHC9534 |
| DMH1 | 1μM | Tocris | 4126 |

**Table 4:** Primary and secondary antibody list.

| **Antibody** | **Host Species** | **Isotype** | **Dilution** | **Brand** | **Catalogue Number** |
| --- | --- | --- | --- | --- | --- |
| **SOX10** | Mouse | IgG<sub>1</sub> | 1:200 | SantaCruz | sc-365692 |
| **HOXC9** | Mouse | IgG<sub>2b</sub> | 1:200 | SantaCruz | sc-81100 |
| **SOX9** | Rabbit | Polyclonal | 1:500 | Sigma | HPA001758 |
| **AlexaFluor555 anti-IgG<sub>1</sub> mouse** | Goat |  | 1:1000 | ThermoFisher Scientific | A-21127 |
| **AlexaFluor488 anti-IgG<sub>2b</sub> mouse** | Goat |  | 1:1000 | ThermoFisher Scientific | A-21141 |
| **AlexaFluor anti-H+L rabbit** | Goat |  | 1:1000 | ThermoFisher Scientific | A-21245 |

**Table 5:**
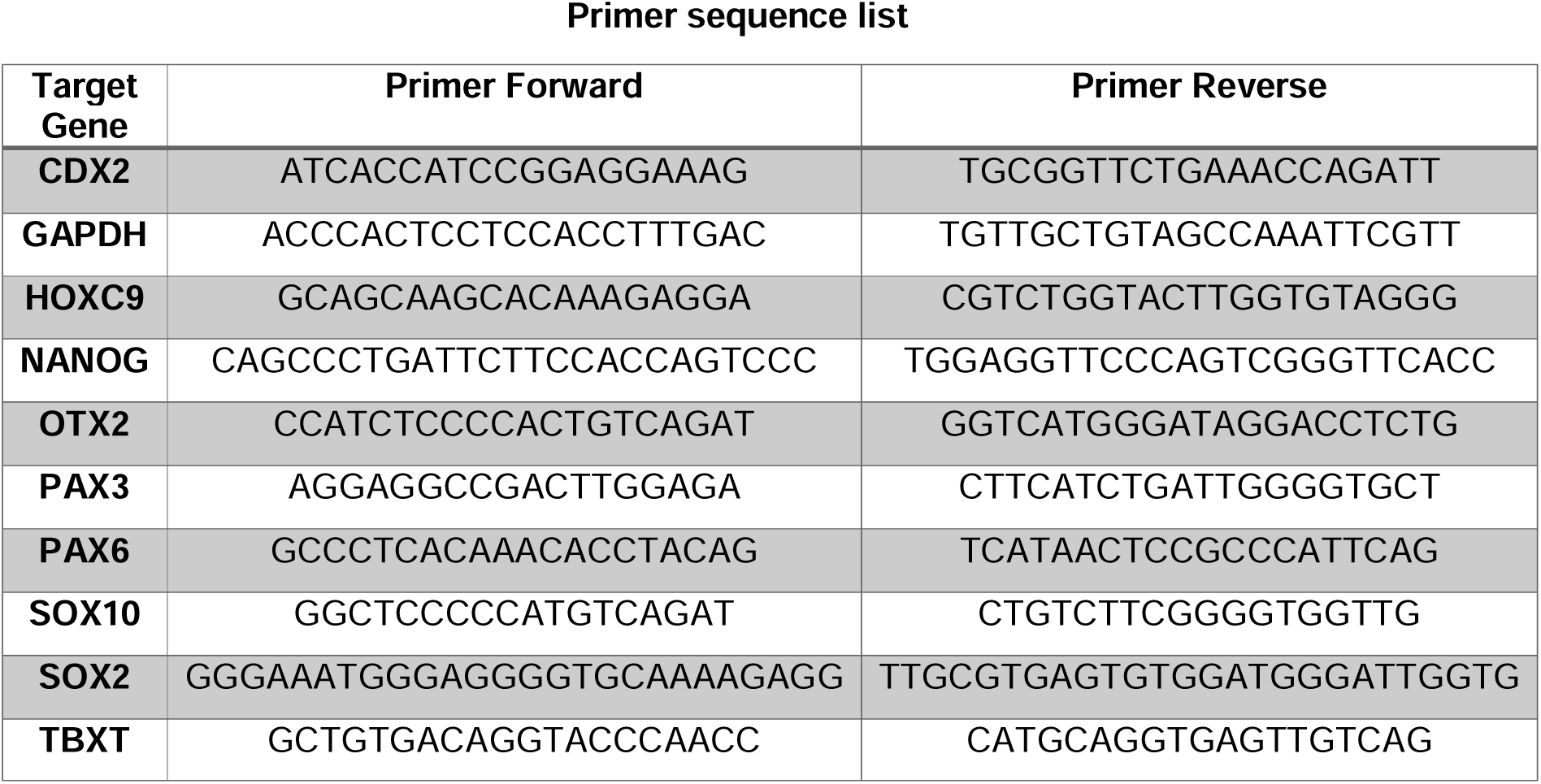
List of primer sequences used in qPCR gene expression analysis.

## Notes

### Competing Interest Statement

The authors have declared no competing interest.

### Funding

- Medical Research Council: MC_PC_21044, MR/W006820/1
- Cancer Research UK: DRCPFA-Nov24/100001
- UK Dementia Research Institute

## References

1. Monsonego-Ornan, E. et al. Matrix metalloproteinase 9/gelatinase B is required for neural crest cell migration. Dev Biol 364, 162–177 (2012).

2. Bronner, M. E. & LeDouarin, N. M. Development and evolution of the neural crest: An overview. Dev. Biol. 366, 2–9 (2012).

3. Theveneau, E. & Mayor, R. Neural Crest Cells. *Part*C*: Neural Crest Cell Evol*. Dev. 73–88 (2014) doi:10.1016/b978-0-12-401730-6.00004-1.

4. Prasad, M. S., Charney, R. M. & García-Castro, M. I. Specification and formation of the neural crest: Perspectives on lineage segregation. Genesis 57, e23276 (2019).

5. Shen, H. et al. Chicken Transcription Factor AP-2: Cloning, Expression and Its Role in Outgrowth of Facial Prominences and Limb Buds. DEVELOPMENTAL BIOLOGY (1997).

6. Luo, T., Lee, Y.-H., Saint-Jeannet, J.-P. & Sargent, T. D. Induction of neural crest in Xenopus by transcription factor AP2α. Proc. Natl. Acad. Sci. 100, 532–537 (2003).

7. Bioethics, N. C. on. Nuffield Council on Bioethics. (2017).

8. Yuan, Y. et al. 3D reconstruction of a human Carnegie stage 9 embryo provides a snapshot of early body plan formation. Cell Stem Cell (2025) doi:10.1016/j.stem.2025.04.007.

9. Dong, R. et al. Single-Cell Characterization of Malignant Phenotypes and Developmental Trajectories of Adrenal Neuroblastoma. Cancer Cell 38, 716–733.e6 (2020).

10. O’Rahilly, R. & Müller, F. The development of the neural crest in the human. J. Anat. 211, 335–351 (2007).

11. Betters, E., Liu, Y., Kjaeldgaard, A., Sundström, E. & García-Castro, M. I. Analysis of early human neural crest development. Dev. Biol. 344, 578–592 (2010).

12. Webb, S. et al. An integrated single cell and spatial omics atlas of human prenatal development. (2026) doi:10.64898/2026.03.30.714220.

13. Vega-Lopez, G. A., Cerrizuela, S., Tribulo, C. & Aybar, M. J. Neurocristopathies: New Insights 150 Years After the Neural Crest Discovery. Dev Biol 444, S110–S143 (2018).

14. Moris, N. et al. An in vitro model of early anteroposterior organization during human development. Nature 582, 410–415 (2020).

15. Hamazaki, N. et al. Retinoic acid induces human gastruloids with posterior embryo-like structures. Nat. Cell Biol. 1–14 (2024) doi:10.1038/s41556-024-01487-8.

16. Gribaudo, S. et al. Self-organizing models of human trunk organogenesis recapitulate spinal cord and spine co-morphogenesis. Nat. Biotechnol. 42, 1243–1253 (2024).

17. Bajpai, R. et al. CHD7 cooperates with PBAF to control multipotent neural crest formation. Nature 463, 958–962 (2010).

18. Prescott, S. L. et al. Enhancer Divergence and cis-Regulatory Evolution in the Human and Chimp Neural Crest. Cell 163, 68–83 (2015).

19. Taroc, E. Z. M. et al. Human ectodermal organoids reveal the cellular origin of DiGeorge Syndrome. bioRxiv 2025.08.08.669417 (2025) doi:10.1101/2025.08.08.669417.

20. Rockel, A. F., Wagner, N., Spenger, P., Ergün, S. & Wörsdörfer, P. Neuro-mesodermal assembloids recapitulate aspects of peripheral nervous system development in vitro. Stem Cell Rep (2023) doi:10.1016/j.stemcr.2023.03.012.

21. Motoike, S., Inada, Y., Toguchida, J., Kajiya, M. & Ikeya, M. Jawbone-like organoids generated from human pluripotent stem cells. *Nat*. Biomed. Eng. 1–19 (2025) doi:10.1038/s41551-025-01419-3.

22. Seto, Y., Ogihara, R., Takizawa, K. & Eiraku, M. In vitro induction of patterned branchial arch-like aggregate from human pluripotent stem cells. Nat. Commun. 15, 1351 (2024).

23. Xue, X. et al. A patterned human neural tube model using microfluidic gradients. Nature 628, 391–399 (2024).

24. Karzbrun, E. et al. Human neural tube morphogenesis in vitro by geometric constraints. Nature 1–5 (2021) doi:10.1038/s41586-021-04026-9.

25. Mitrofanova, O. et al. Bioengineered human colon organoids with in vivo-like cellular complexity and function. Cell Stem Cell 31, 1175–1186.e7 (2024).

26. Hagemann, C. et al. Low-cost, versatile, and highly reproducible microfabrication pipeline to generate 3D-printed customised cell culture devices with complex designs. PLOS Biol. 22, e3002503 (2024).

27. Abe, Y., Fukai, D., Toyoda, T., Fukumoto, N. & Terao, K. PVA-coated 3D-printed molds for rapid prototyping of PDMS microdevices for stem cell culture. Biomed. Microdevices 28, 10 (2026).

28. Shaker, M. R. et al. Spatiotemporal contribution of neuromesodermal progenitor-derived neural cells in the elongation of developing mouse spinal cord. Life Sci. 282, 119393 (2021).

29. Frith, T. et al. Human axial progenitors generate trunk neural crest cells in vitro. Elife 7, e35786 (2018).

30. Saldana-Guerrero, I. M. et al. A human neural crest model reveals the developmental impact of neuroblastoma-associated chromosomal aberrations. Nat. Commun. 15, 3745 (2024).

31. Hackland, J. O. S. et al. FGF Modulates the Axial Identity of Trunk hPSC-Derived Neural Crest but Not the Cranial-Trunk Decision. Stem Cell Rep 12, 920–933 (2019).

32. Hackland, J. O. S. et al. Top-Down Inhibition of BMP Signaling Enables Robust Induction of hPSCs Into Neural Crest in Fully Defined, Xeno-free Conditions. Stem Cell Rep 9, 1043–1052 (2017).

33. Garcı a-Castro, M. I., Marcelle, C. & Bronner-Fraser, M. Ectodermal Wnt Function as a Neural Crest Inducer. Science 297, 848–851 (2002).

34. Gomez, G. A. et al. Human neural crest induction by temporal modulation of WNT activation. Dev. Biol. 449, 99–106 (2019).

35. Leung, A. W. et al. WNT/β-catenin signaling mediates human neural crest induction via a pre-neural border intermediate. Development 143, 398–410 (2016).

36. Gomez, G. A. et al. WNT/β-catenin modulates the axial identity of embryonic stem cell-derived human neural crest. Development 146, dev175604 (2019).

37. Ji, Y., Hao, H., Reynolds, K., McMahon, M. & Zhou, C. J. Wnt Signaling in Neural Crest Ontogenesis and Oncogenesis. Cells 8, 1173 (2019).

38. Ray, A. T. et al. FGF signaling regulates development by processes beyond canonical pathways. Genes Dev. 34, 1735–1752 (2020).

39. Creuzet, S., Schuler, B., Couly, G. & Douarin, N. M. L. Reciprocal relationships between Fgf8 and neural crest cells in facial and forebrain development. Proc. Natl. Acad. Sci. 101, 4843–4847 (2004).

40. Bond, A. M., Bhalala, O. G. & Kessler, J. A. The dynamic role of bone morphogenetic proteins in neural stem cell fate and maturation. Dev. Neurobiol. 72, 1068–1084 (2012).

41. Rothstein, M., Bhattacharya, D. & Simoes-Costa, M. The molecular basis of neural crest axial identity. Dev Biol 444, S170–S180 (2018).

42. Taneyhill, L. A. To adhere or not to adhere. Cell Adhes. Migr. 2, 223–230 (2008).

43. Zhao, R. & Trainor, P. A. Epithelial to mesenchymal transition during mammalian neural crest cell delamination. Semin. Cell Dev. Biol. 138, 54–67 (2022).

44. Aoki, Y. et al. Sox10 regulates the development of neural crest-derived melanocytes in Xenopus. Dev. Biol. 259, 19–33 (2003).

45. Cheung, M. & Briscoe, J. Neural crest development is regulated by the transcription factor Sox9. Development 130, 5681–5693 (2003).

46. Friedl, P., Wolf, K. & Lammerding, J. Nuclear mechanics during cell migration. Curr. Opin. Cell Biol. 23, 55–64 (2011).

47. Pérez-Venteo, A., Bosch-Calvet, M., Garcia-Cajide, M. & Mauvezin, C. The responsive nucleus: morphological signatures of cellular state. Nucleus 17, 2630145 (2026).

48. Malagon, S. G. G. et al. Glycogen synthase kinase 3 controls migration of the neural crest lineage in mouse and Xenopus. Nat Commun 9, 1126 (2018).

49. Shellard, A. & Mayor, R. Integrating chemical and mechanical signals in neural crest cell migration. Curr. Opin. Genet. Dev. 57, 16–24 (2019).

50. Theveneau, E. & Mayor, R. Neural crest delamination and migration: From epithelium-to-mesenchyme transition to collective cell migration. Dev. Biol. 366, 34–54 (2012).

51. Szabó, A. & Mayor, R. Mechanisms of Neural Crest Migration. Annu. Rev. Genet. 52, 43–63 (2018).

52. Hagemann, C. et al. Axonal Length Determines Distinct Homeostatic Phenotypes in Human iPSC Derived Motor Neurons on a Bioengineered Platform. Adv Healthc Mater e2101817 (2022) doi:10.1002/adhm.202101817.

53. Cooper, F. et al. Rostrocaudal patterning and neural crest differentiation of human pre-neural spinal cord progenitors in vitro. Stem Cell Rep 17, 894–910 (2022).

54. Paka, C. A., Barrell, W. B., Monsoro-Burq, A. H. & Liu, K. J. Differentiation of hiPSCs to the neural crest lineage (Chapter 6). Advances in Stem Cell Biology (2021).

55. Kirino, K., Nakahata, T., Taguchi, T. & Saito, M. K. Efficient derivation of sympathetic neurons from human pluripotent stem cells with a defined condition. Sci. Rep. 8, 12865 (2018).

56. Yardley, N. & García-Castro, M. I. FGF signaling transforms non-neural ectoderm into neural crest. Dev. Biol. 372, 166–177 (2012).

57. Patthey, C., Edlund, T. & Gunhaga, L. Wnt-regulated temporal control of BMP exposure directs the choice between neural plate border and epidermal fate. Development 136, 73–83 (2009).

58. Glass, M. R. et al. Cross-site reproducibility of human cortical organoids reveals consistent cell type composition and architecture. Stem Cell Rep. 19, 1351–1367 (2024).

59. Hofer, M. & Lutolf, M. P. Engineering organoids. Nat. Rev. Mater. 6, 402–420 (2021).

60. Fattahi, F. et al. DERIVING HUMAN ENS LINEAGES FOR CELL THERAPY AND DRUG DISCOVERY IN HIRSCHSPRUNG’S DISEASE. Nature 531, 105–109 (2016).

61. Danielian, P. S., Muccino, D., Rowitch, D. H., Michael, S. K. & McMahon, A. P. Modification of gene activity in mouse embryos in utero by a tamoxifen-inducible form of Cre recombinase. Curr. Biol. 8, 1323–S2 (1998).

62. Muzumdar, M. D., Tasic, B., Miyamichi, K., Li, L. & Luo, L. A global double-fluorescent Cre reporter mouse. *genesis* **45**, 593–605 (2007).

63. Choi, H. M. T. et al. Mapping a multiplexed zoo of mRNA expression. Development 143, 3632–3637 (2016).

