## Supplementary figures and images for "Dynamic modelling of human neural crest development using a bioengineered stem cell organoid system"

### Supplemental Figure 1

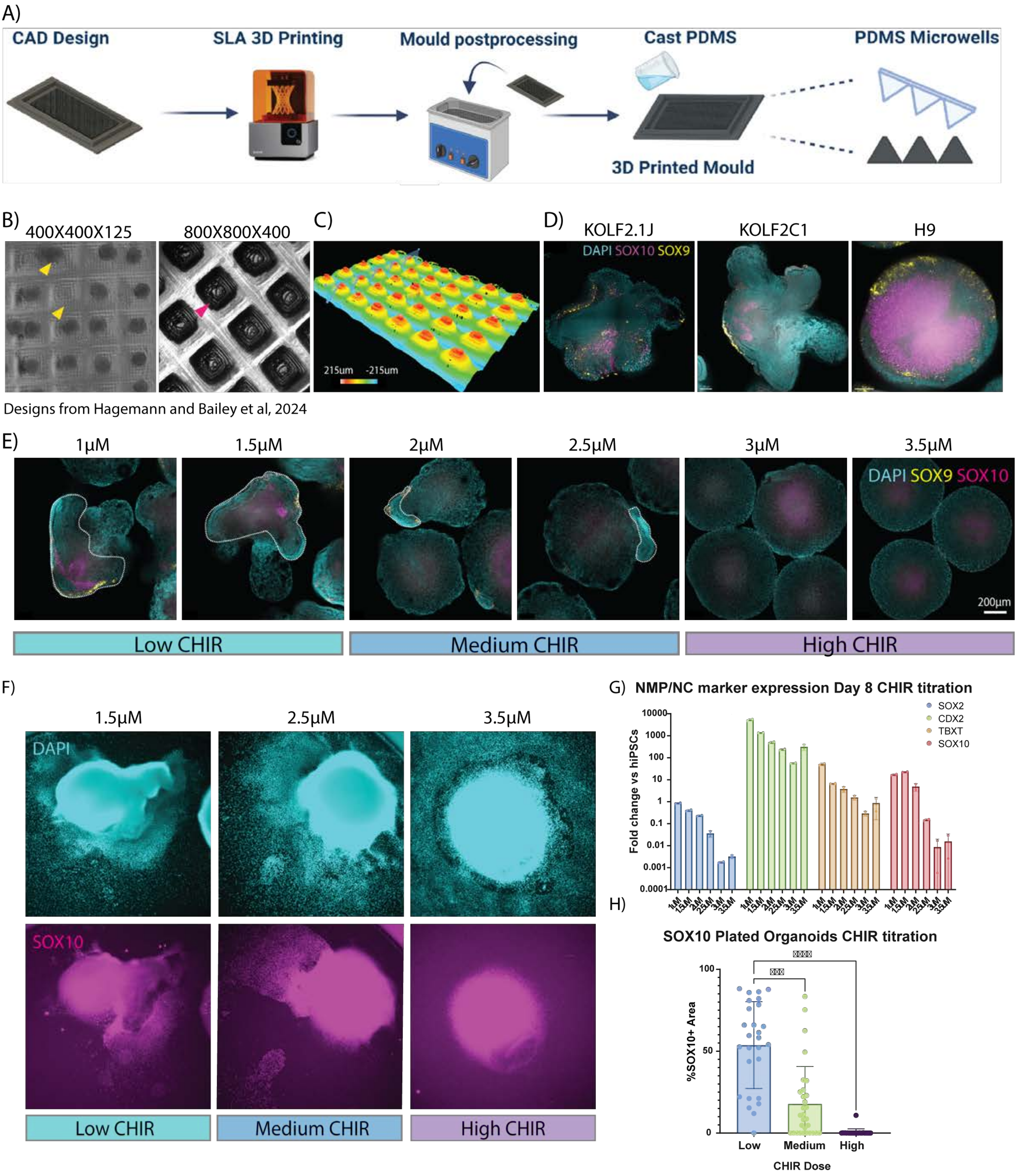

### Supplemental Figure 2

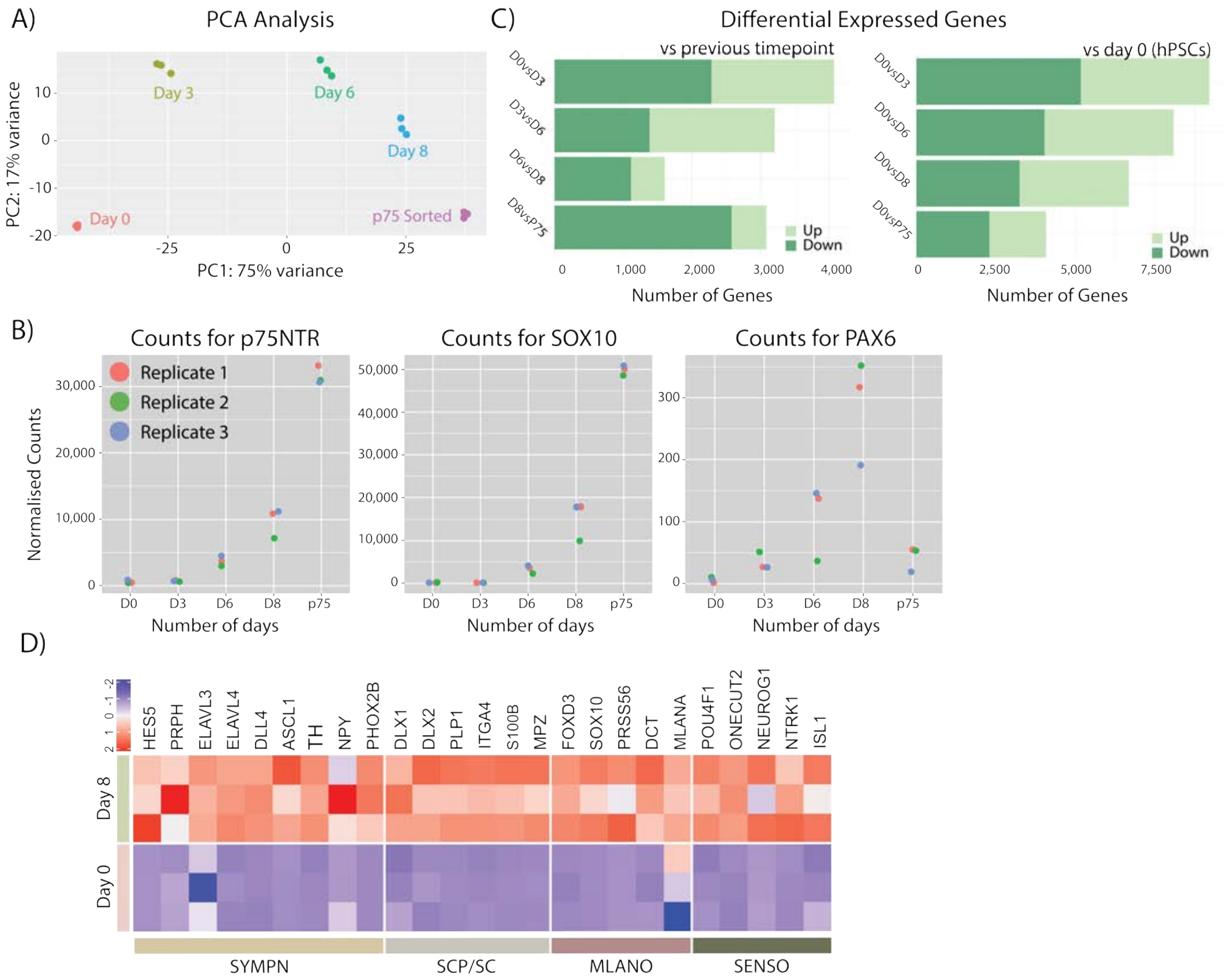

### Supplemental Figure 3

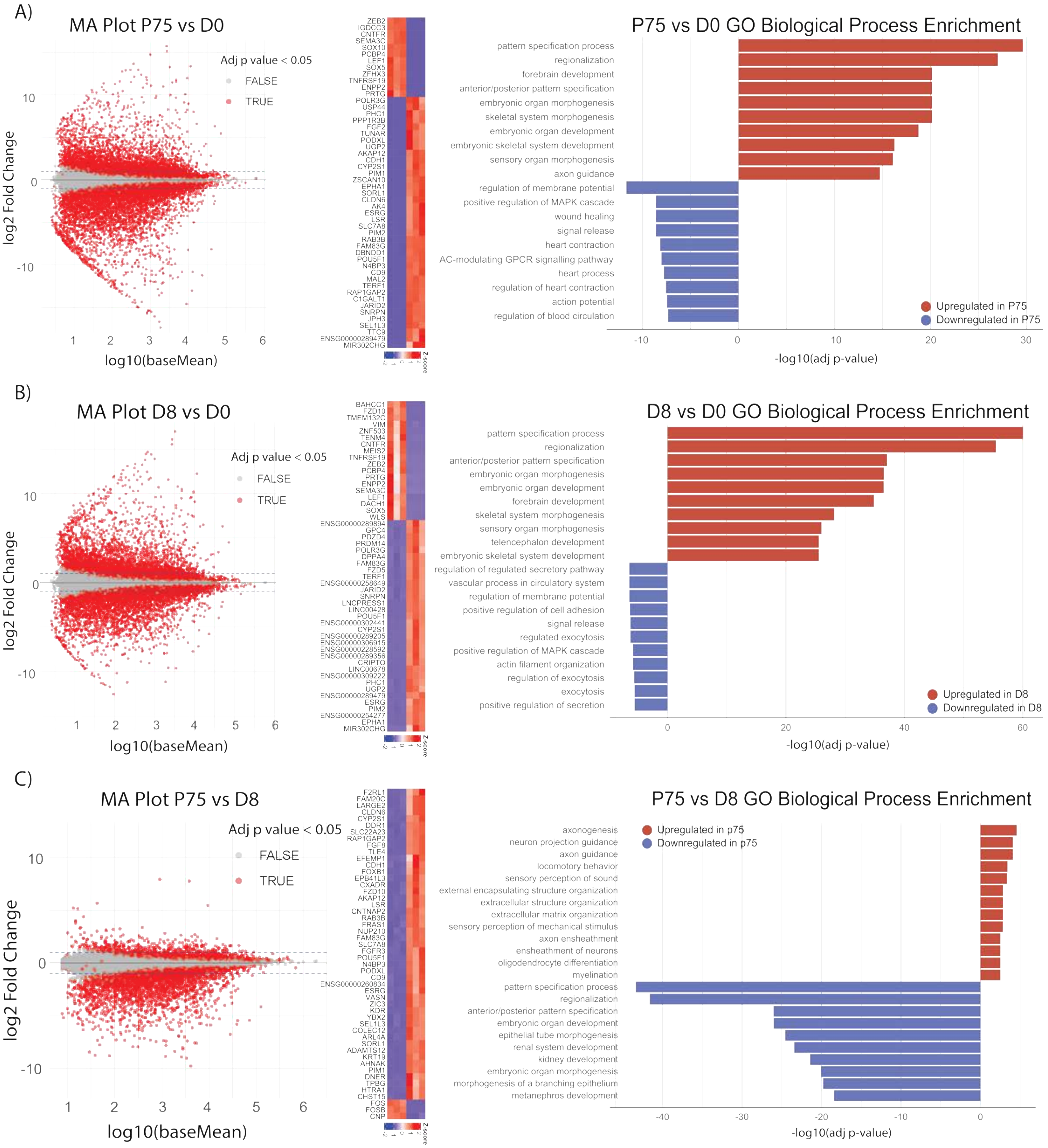

### Supplemental Figure 4

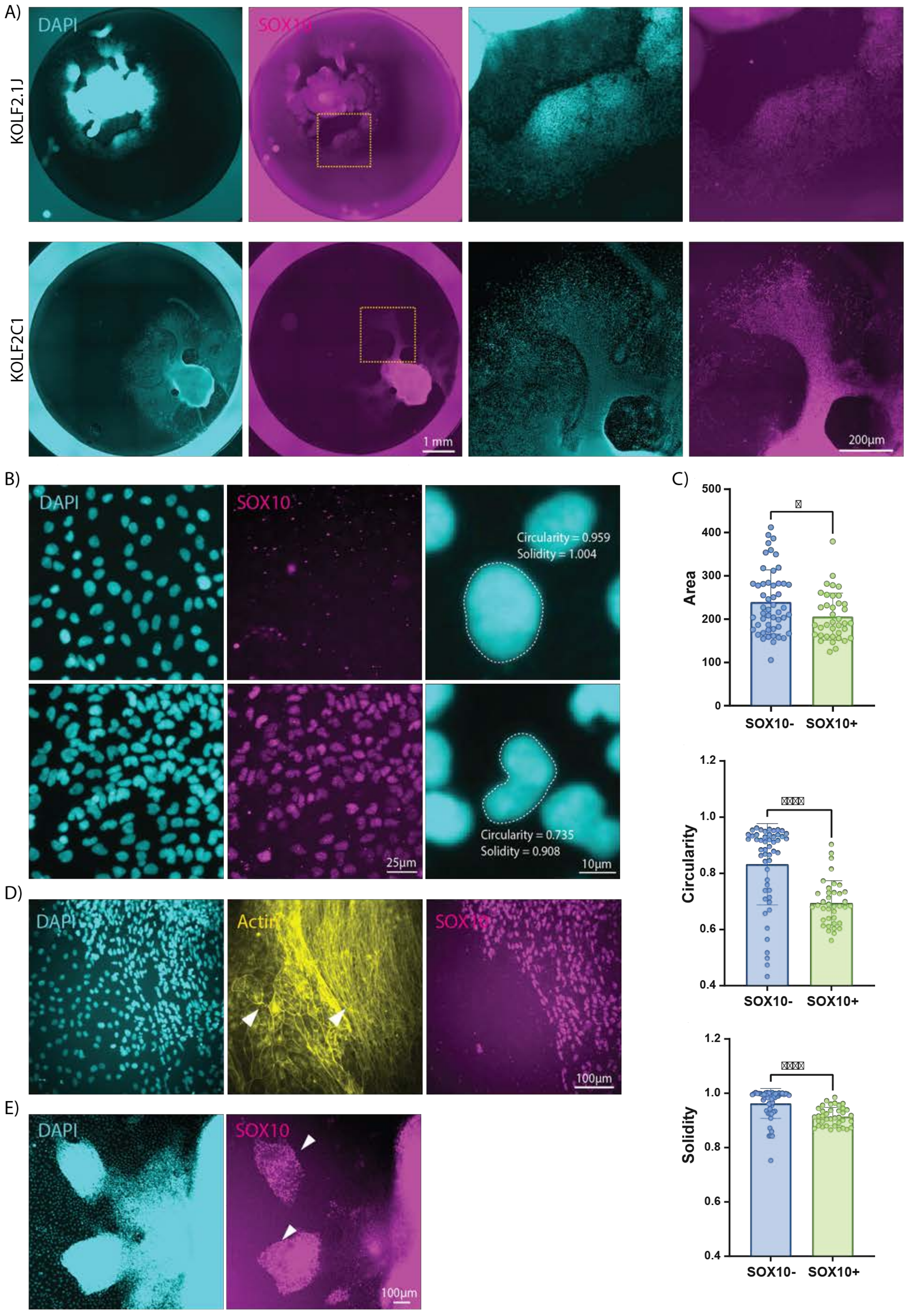

### Supplemental Figure 5

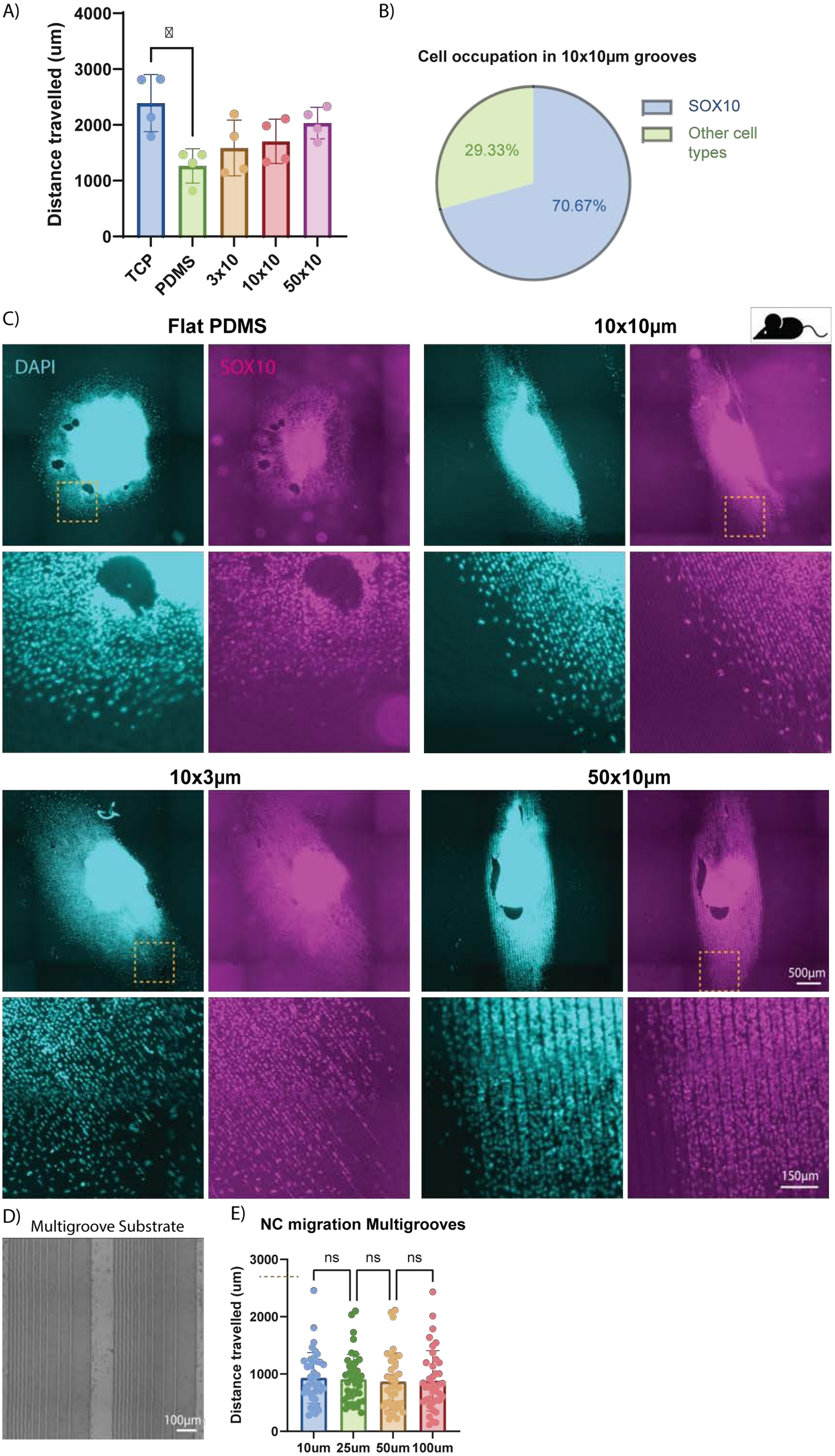
